# Impact of commercial gut health interventions on caecal metagenome and broiler performance

**DOI:** 10.1101/2024.08.02.606333

**Authors:** Gladys Maria Pangga, Banaz Star-Shirko, Androniki Psifidi, Dong Xia, Nicolae Corcionivoschi, Carmel Kelly, Callie Hughes, Ursula Lavery, Anne Richmond, Umer Zeeshan Ijaz, Ozan Gundogdu

**Author notes:** Corresponding authors: Ozan Gundogdu, Assistant Professor, London School of Hygiene and Tropical Medicine; Umer Zeeshan Ijaz, James Watt School of Engineering, University of Glasgow, Glasgow, UK.

## Abstract

**Background:** Maintaining gut health is a persistent and unresolved challenge in the poultry industry. Given the critical role of gut health in chicken performance and welfare, there is a pressing need to identify effective gut health intervention (GHI) strategies to ensure optimal outcomes in poultry farming. In this study, across three broiler production cycles, we compared the metagenomes and performance of broilers provided with ionophores as control against birds subjected to five different GHI combinations involving vaccination, probiotics, prebiotics, essential oils, and reduction of ionophore use.

**Results:** Using a binning strategy, 84 (≥75% completeness, ≤ 5% contamination) metagenome-assembled genomes (MAGs) from 118 caecal samples were recovered and annotated for their metabolic potential. The majority of these (n = 52, 61%) had a differential response across all cohorts and are associated with the performance parameter - European Poultry Efficiency Factor (EPEF). The control group exhibited the highest EPEF, followed closely by the cohort where probiotics are used in conjunction with vaccination. The use of Probiotics B, a commercial Bacillus strain-based formulation, was determined to contribute to the superior performance of birds. GHI supplementation generally affected abundance of microbial enzymes relating to carbohydrate and protein digestion, and metabolic pathways relating to energy, nucleotide synthesis, short-chain fatty acid synthesis, and drug-transport systems. These shifts are hypothesised to differentiate performance among groups and cycles, highlighting the beneficial role of several bacteria including *Rikenella microfusus* and UBA7160 species.

**Conclusions:** All GHIs are shown to be effective methods for gut microbial modulation, with varying influences on MAG diversity, composition and microbial functions. These metagenomic insights greatly enhance our understanding of microbiota-related metabolic pathways, enabling us to devise strategies against enteric pathogens related to poultry products and presenting new opportunities to improve overall poultry performance and health.

## Introduction

The gastrointestinal tract of chickens harbours a complex and dynamic microbial community collectively known as the gut microbiota. This microbiota, along with its corresponding genetic material, forms the gut microbiome which is recognised for its significance in both health and metabolism in its host [1]. The majority of the microbiome consists of a diverse set of bacteria which can be classified as either commensal, pathogenic or beneficial to the host; of which their occurrence and interactions can be influenced by a variety of factors, such as genetics, age, environment, diet, and administration of feed additives [1–4]. Interestingly, these same factors have also been established to directly impact overall health and performance of the chicken host [5–7], indicating a possible link between gut microbiota composition and broiler performance. Indeed, one study identified 24 bacterial species to be differentially abundant between broilers with high and low feed conversion ratios (FCR) [8]. Furthermore, in our previous work, we demonstrated that extrinsic parameters including stocking density, percentage of protein and energy in diet, and Omega-3 supplementation are able to modulate key microbiome members involved in nutrition and metabolism, subsequently affecting growth and feed efficiency in the host [9].

Due to this accumulating evidence supporting the importance of gut health in poultry performance, there has been a significant rise in interest on modulation of gut microbiota for improved animal health, productivity, and food safety [10]. Historically, growth promoters have utilised for enhanced feed efficiency, while also decreasing illness and death rates from both overt and hidden diseases; hence, [11]. These drugs are purported to achieve these benefits by altering the gut microbiota, resulting in decreased nutrient utilisation by microbes, increased absorption of nutrients through thinned host gut walls, and reduction in inflammatory stress [11–13]. Ionophores are growth promoters that have been safely used as the approved and standard intervention to maintain gut health in poultry in the last decades, especially for the effective prevention of economically important diseases such as coccidiosis and necrotic enteritis [14–16]. Currently, a myriad of new gut health interventions (GHIs) with similar effects are being considered as supplements to further improve overall bird health and performance. These include, but are not limited to prebiotics, probiotics, phytogenic substances, organic acids, essential oils, and enzymes [17]. Each of these GHIs has their own mechanism of action and corresponding effects on chickens, with different subtypes within each group. Descriptions of these GHIs have been detailed previously in recent reviews [18,19]. Options are further expanded by combinatory use of multiple GHIs throughout a single production cycle for their potential synergistic effects [20,21]. However, there is limited information on their effects on the gut microbiome in poultry.

Research in order to investigate the influence of the GHIs has mainly involved the use of metataxonomic sequencing of the microbiota through amplification of the 16S rRNA gene marker [22]. Use of this method has proved ground-breaking for our understanding of gut health; however, taxa identification of less abundant and unknown species as well as the characterising of their metabolic capacity (functional profiles) remains challenging [22–24]. These limitations can hinder our comprehension of their connection to broiler health and performance. In contrast, with shotgun sequencing (which involves indiscriminate sequencing of all random DNA segments within a sample), a higher resolution of microbial genomes enables these features to be identified, allowing a deeper understanding of relevant metabolic functions of bacteria [24]. For instance, Chen et al., (2023) [25] used metagenomic methods in order to understand the role of gut microbiota in fat regulation in chickens. From this, they were able to observe the presence of differential carbohydrate active enzymes (CAZymes) and functional metabolic modules between high and low abdominal fat chickens. Applying this concept, an exploration of the metagenome of chickens given various GHIs will help us understand the gut microbial functions of chickens that promote an improved health and performance of broilers. Therefore, the aim of this study is to characterise how GHIs impact the gut microbiome in relation to performance through shotgun metagenomic sequencing, in comparison to the standard use of ionophores.

## Materials and methods

### Ethics statement

All animal trials were reviewed and conducted in accordance with the Animal Welfare and Ethical Review Board of London School of Hygiene and Tropical Medicine, UK (Reference 2023-05). Poultry farm management and industry plant processing activities were conducted following Moy Park Ltd standard operating protocols which are compliant with UK animal handling laws and regulations (Craigavon, UK) (Moy Park Ltd, 2022, 2023) [26–27]. As part of standard commercial practices of the company, all birds were subjected to stunning before slaughter and subsequent carcass processing [28].

### Experimental design and sampling

Three broiler production cycles (C0, C1, C2) were implemented in a Moy Park Ltd affiliated commercial farm in Northern Ireland between June and October 2022. In each cycle, a total of 18,000 Ross-308 mix-sexed broilers were raised in an automated commercial poultry house and provided a four-stage standard commercial diet regimen based on Aviagen specifications for Ross broilers [29]. This was composed of a starter diet (S, 0-11 days), grower diet (G, 11-23 days), finisher diet (F, 23-31 days) and a withdrawal diet (W, 32 days until clearing at day 40). Birds were distributed into 6 groups (1 control, 5 GHI groups), wherein each group is allocated into 6 pens containing 500 birds each. For our control group, we administered ionophores which is a safe and legally accepted method for control of coccidiosis [15,16]. For our treatment groups, we adapted different gut health strategies which involves combination of GHIs; this was designed to optimise and maximise the differences on gut health and performance as based on the study of Granstad et al., (2020). GHI treatment groups for C0 and C1 includes: T2 – Coccidiosis vaccine (V); T3 – V + *Bacillus* strain probiotics A (PA), T4 – V + PA + reduced crude protein (-1%) in G/F/W diets; T5 – V + PA + Essential oil and T6 – V + PA + and ionophore in F diet. Birds in C2 had similar treatment design with two differences: a different GHI – *Bacillus* strain Probiotics B (PB) was utilised instead of PA in T2 to T5, and essential oils in T5 was replaced by prebiotics. Birds were provided *ad libitum* access to feed and water.

At the end of the production cycle (Day 40), caecal samples were collected within 5-10 minutes of the slaughtering process in a Moy Park Ltd industry plant. Intact caeca (N = 120) from C1 (n = 60) and C2 (n = 60) were obtained through aseptic incision from the rest of the GIT and were then transferred into sterile 50ml tubes and stored in a polystyrene container with frozen icepacks. All specimens were immediately sent to the laboratory for storage at -80°C until further use for DNA extraction (Illustrated in Figure 1).

**Figure 1.**
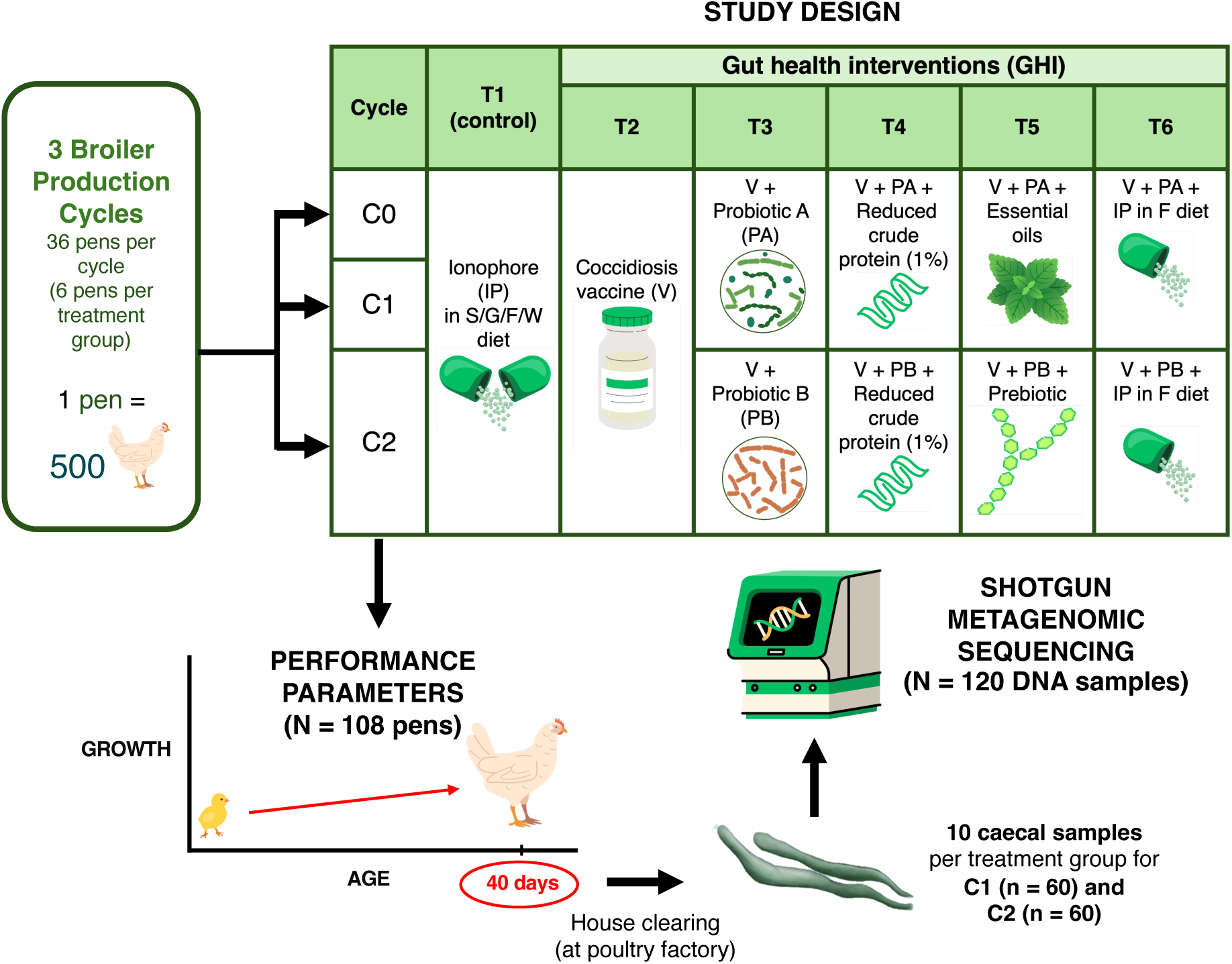
Overview of study design. C: Production cycle, S: Starter, G: Grower, F: Finisher, W: Withdrawal, PA: Probiotic A, PB: Probiotic B, IP: Ionophore, V: Coccidiosis vaccine

### DNA extraction, library preparation, and shotgun metagenomic sequencing

Microbial DNA was extracted from caecal chyme, using the QIAamp® PowerFecal Pro DNA Kit (Qiagen, Germany) according to manufacturer’s instructions. Total DNA was eluted in 50 µl of elution buffer and stored at -80°C. Initial DNA concentration was measured using Nanodrop ND-1000 (NanoDrop Technologies, Inc., Wilmington, US).

Sequencing libraries were generated using a modified Illumina DNA Prep tagmentation approach (Illumina, Inc., Cambridge, UK) described previously [30]. Tagmentation was performed as follows: a master mix composed of 0.5 µl bead linked transposomes 0.5 µl tagmentation buffer and 4 µl nuclease free distilled water was created for each sample (2 µl), placed in a 96-well plate, and run on a thermocycler at 55°C for 15 mins. Another PCR master mix using the Kapa2G Fast Hot Start PCR kit (Sigma-Aldrich, Gillingham, UK) was then generated and transferred into a 96-well plate, to which 5 µl of P7 and P7 of Nextera XT Index Kit v2 index primers (Illumina, Cambridge, UK) and 7 µl of the previous tagmentation reaction were added. The plate was run on the thermocycler with conditions: 72°C for 3 mins, 95°C for 1 min, 14 cycles of 95°C for 10 secs, 55°C for 20 secs and 72°C for 3 mins. Quality control of multiplex barcoding was performed on a D5000 ScreenTape using the Agilent Tapestation 4200 (Agilent, Wardbronn, Germany). Next, barcoded libraries were quantified on a Qubit 3.0 instrument (Invitrogen, Paisley, UK), pooled in equivalent concentrations in a tube, and washed with 0.5-0.7X solid phase reversible immobilisation KAPA Pure Beads (Roche, Wilmington, US). In order to calculate final pool molarity, the pooled library was quantified using Qubit 3.0 and on a D5000 High Sensitive ScreenTape. After library qualification, the library was sequenced using the NovaSeq 6000 System, Paired-end 150 bp (Illumina, Cambridge, UK).

### Bird performance and health monitoring

The performance parameters included bird weight (BW, kg of body weight), average daily gain (ADG, grams feed per day), corrected food conversion ratio (FCR) at 2kg BW, and total mortality (MT). These measurements were taken as mean average per pen at clearing day (40d) which were conducted in line with typical industrial practices. Contact dermatitis measures which included footpad dermatitis (FPD) lesion scores (FPDS), FPD prevalence (FPDP), and hockburn (HB) lesion scores (HBS) and prevalence (HBP) were also taken before slaughter as conducted previously [5]. To estimate overall performance, European Production Efficiency Factor (EPEF) was calculated based on preliminary recorded performance measures [31].

## Bioinformatic analysis

### Recovery of Metagenomic-Assembled Genomes (MAGs)

A total of 120 metagenomic samples were processed – from which adapter trimmed reads were generated by the sequencing centre. Reads were subjected to quality trimming using Sickle v1.200 [32]. Trimming involved removing reads where the average phred below 20 and retaining paired end reads with a post-trimming length exceeding 50 bp. Two samples (one from T3 and one from T4 in C2) were excluded due to non-recovery of reads, resulting in a total of 118 samples which generated 2,588,938,595 reads. Forward and reverse reads were then aggregated and subjected to collective assembly using MEGAHIT [33]. Assembly parameters used were --k-list 27,47,67,87 --kmin-1pass -m 0.95 --min-contig-len 1000. This gave us a total of 1,276,325 contigs, a total of 3,101,580,806 bases (bp), maximum of 403,439 bp, average length of 2,430 bp, and an N50 score of 2,724 bp. Assemblies were then subjected to binning via the MetaWRAP pipeline [34], wherein three algorithms namely metabat2 [35], MaxBin [36], and CONCOCT [37] were utilised. Bins from each of the algorithms were consolidated using the MetaWRAP framework, resulting in a total of 308 bins. For estimation of completion (COM) and contamination (CON) metrics of each MAG, CheckM was used on all bins [38]. We retained bins with more than 75% and less than 5% contamination to give a final set of 84 MAGs. The summary statistics of these MAGs are provided in Supplement file 1.

### Taxonomic and functional annotation

For metabolic function and taxonomic assessment of each MAG, the METABOLIC pipeline was employed [39]. Within its framework, taxonomic classification of bins was incorporated using GTDB-TK [40], whilst functional annotations were recovered using Kyoto Encyclopedia of Genes and Genomes (KEGG) for metabolic function modules and submodules [41], dbCAN2 for carbohydrate active enzymes (CAZymes) [42], custom hidden Markov model databases for nutrient cycles [43] and MEROPS for proteases [44]. To obtain taxonomic and functional coverages per sample, read coverages (proportion of each bin per sample) were multiplied with each feature coverages (returned from METABOLIC). From this, we derived the sample-wise abundance tables: dbCAn2 (n = 118 samples x 117 CAZyme IDs), KEGG Modules (n = 118 samples x 251 module IDs), KEGG Submodules (n = 118 samples x 964 submodule IDs) and MEROPS (n = 118 samples x 108 peptidases).

### Phylogenetic tree generation

To construct a phylogenetic tree of MAGs, we used GToTree [45] that involves detection of Single Copy Genes (SCGs) in MAGs and multisequence alignment. Specifically, we used the bacteria and archaea HMM set which covers 25 SCGs. MAGs that had very few hits for these SCG were removed, resulting in a phylogeny recovery for a total of 65 MAGs. For assessment of novelty of MAGs, the Genome Tree Toolkit was utilised [46], wherein phylogenetic gain (PG) for each MAG against other MAGs in the tree was estimated.

#### Statistical analyses

All tests were performed in R [47]. The normality of data was assessed using Shapiro-Wilk test [48]. To determine significant differences between treatment groups, we employed analysis of variance (ANOVA) and pairwise t-test with Bonferroni correction for normally distributed data, while Kruskal-Wallis testing and *posthoc* Dunn testing with Bonferroni correction (p < 0.05) for non-normal distributed data [47]. For performance data, when significant p-values (p < 0.05) were obtained in ANOVA, statistical groupings were evaluated using Duncan Multiple Range Test (DMRT) through the *agricolae* package [49].

To evaluate the individual effects of GHIs across the three cycles, we performed a generalised linear model analysis via the penalised maximum likelihood method using the *glmnet* package [50]. Specifically, we used the least absolute shrinkage and selection operator (LASSO) or L1 penalty model by setting the regularisation parameter or *alpha* = 1 and cross validation folds (*nfolds*) = 10. This generalised model was used to prevent overfitting of models due to multicollinearity and sparsity of covariates (such as vaccination and ionophore use in our study) and to determine which covariate fits best as a predictor of the outcome of interest. The model is represented as:

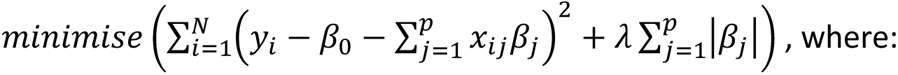

𝑁: the number of observations, 𝑝: the number of predictors, 𝑦_i_: the outcome variable for the 𝑖-th observation, 𝑥_ij_: the value of the 𝑗 -th predictor for the 𝑖-th observation, 𝛽_0_: intercept term, 𝛽_j_: are the coefficients for the parameter, and 𝜆: the regularisation parameter controlling the strength of the penalty term [51]. This model indicates that the penalty term forces some of the beta coefficients to go to zero when their corresponding predictors are not significant. For our model, we included the following as predictors: Ionophore: All stages (as “Yes”) or “No”, Ionophore: Finisher (as “Yes”) only or “No”, Vaccination: “Yes” or “No”, Probiotic A: “Yes” or “No”, Probiotic B: “Yes” or “No”, Essential oil: “Yes” or “No”, Reduced crude protein (-1%): “Yes” or “No”, Prebiotic: “Yes” or “No”, wherein “No” was used as reference for all covariates. For outcome variables, we used the parameters: 40d ADG, 40d BW, EPEF, corrected 2kg FCR (2kgFCR), FPDP (%), HBP (%), and total MT (%).

For microbial diversity assessment, different functions of the *vegan* package [52] were employed. For alpha diversity, we estimated richness (R) (using the *rarefy* function), Shannon entropy (H) and Simpson (Si) (using the *diversity* function), Fisher alpha (FA) (using the *fisher.alpha* function), and Pielou’s evenness (PE) (using the *specnumber* (S) function for formula: 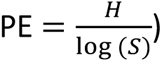. After confirming their normal distribution, ANOVA was then employed to determine significance differences between treatment groups [47]. For beta diversity, we employed Bray Curtis dissimilarity index analysis using the *vegdist* function of the *vegan* package, followed by principal component analysis using the R base function *cmdscale* [47,52]. The separation between groups was tested with permutational analysis of variance (PERMANOVA) through the *vegan* command *adonis2* [52].

To find the relationship between the individual MAGs and each treatment group (C1 and C2 done separately) as well as the relationship between individual MAGs and performance parameters – EPEF and MT, we employed Generalised Linear Latent Variable Model (GLLVM) regression analysis as described previously [53]. This model is an extension of the basic generalised linear model wherein the mean abundances (for *i-*th sample and *j-*th MAG) are regressed against the covariates (***T,*** treatment groups or performance/health trait) by incorporating latent variables 𝒖_𝒊_ as:

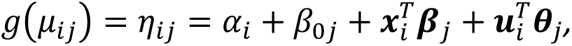

where **𝛽***_j_* are coefficients specific to each MAGs associated with individual covariate, 𝜽*_j_* are the coefficients associated with the latent variable, and 𝛽*_0j_* are MAG-specific intercepts, while 𝛼*_i_* are optional sample effects (which can either be random or fixed effects) and 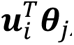, are random effects (latent variables). The recovered **𝛽***_j_* coefficients form a relationship between the covariate of interest *j* and the abundance of a microbe *i* with a positive value suggesting an increase in the covariate causes an increase in abundance whilst the negative value shows a converse response. The 95% confidence intervals of these coefficients were generated and where they crossed the 0 boundary, they were deemed insignificant. We conducted this GLLVM analysis through the *gllvm* package, where we specified use of the negative binomial distribution and the variational approximation method [54].

For comparison of grouped features between treatment groups, sample-wise abundance tables were initially subjected into normalisation by total sum scaling (TSS) (number of reads for each MAG divided by total of reads per sample) and subsequently via centralised log ratio (CLR) method using the *logratio.transfo* function (log-ratio transformation) of the *mixOmics* package [55]. Meanwhile, for individual feature statistical comparison between treatment groups, sample-wise abundance tables were subjected into differential expression analysis using the *DESeq2* package with default settings (test: Negative binomial Wald test, type of fitting of dispersions to the mean intensity: *parametric*, with p < 0.05) [56]. Individual CAZyme function categorisation was adapted from dbCAN2 annotated substrate information [42] and ontology of MetaCyc database [57], with supplementing information from the CAZy database [58] and CAZypedia [59]. Identification of enzymes and subsequent KO enzyme mapping for production pathways of short-chain fatty acids (SCFA): acetate, butyrate and propionate were adapted from previous metagenomic studies [60,61]. Identified enzymes were then matched to KEGG submodules using the KEGG database [41].

For visualisation, *ggplot2* was used for generation of plots (line graphs, barplots, boxplots, sankeyplot) [62], while *ComplexHeatMap* was utilised for heatmap clustering (we used Euclidean clustering) [63]. For mapping of phylogenetic tree, we utilised the packages *ape* and *ggtree* for manipulation and layering of other MAG features: Guanosine-Cytosine (GC) content, novelty (PG), and quality score (computed using formula: COM – 5 x CON) [64,65].

## Results

### Administration of different GHIs impacted overall bird performance

Growth performance is one of the most important indicators of nutrition and health in poultry. According to Table 1, significant differences between treatment groups were only observed in FCR, EPEF, and HB measures (ANOVA, p < 0.05). Estimation of corrected FCR of C0 and C2 has shown T1 to have best FCR. EPEF was observed to be highest in T1, followed by T3, but was not significant in C0 and C1. Across all cycles, performance of C2 can be considered most superior as EPEF, BW and ADG of C2 are consistently higher than the C0 and C1 (ANOVA, p <0.05, Table 2). However, C2 also demonstrated the highest values and significant between-treatment differences in HB scores and prevalence, in which T1 and T2 exhibited lowest, whilst T4 exhibited the highest HB metrics (ANOVA, p < 0.05).

**Table 1.**
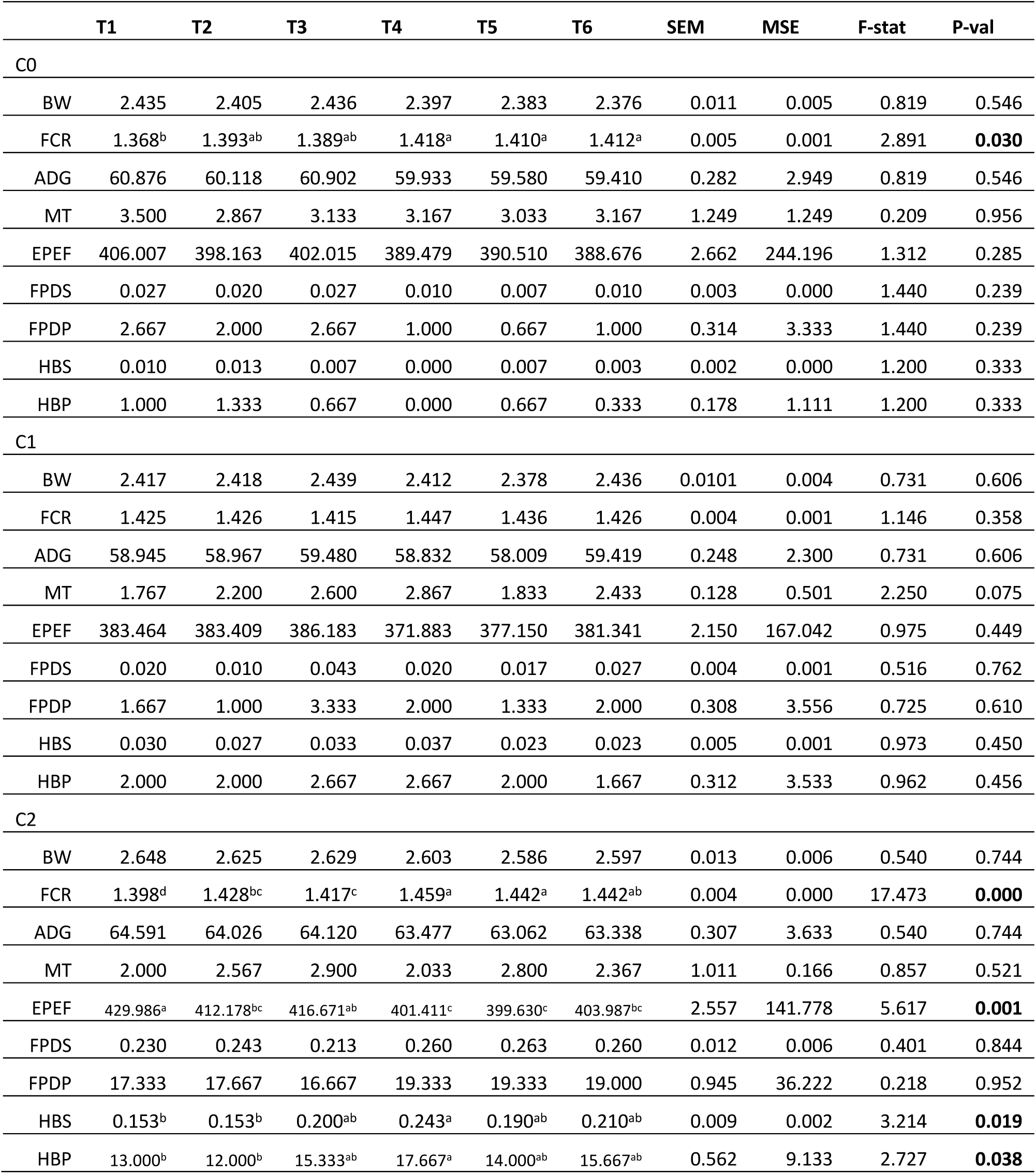
Performance and health parameters of birds across treatment groups grouped according to cycles. Cycle 0 (C0), Cycle 1 (C1) and Cycle 2 (C2) (N = 108 pens, 36 pens/cycle). Performance parameters taken as mean average per pen: Bird weight (BW), Average daily gain (ADG) - grams per day, Feed conversion ratio (FCR), Total mortality (MT), (E) European Production Efficiency Factor (EPEF), Footpad dermatitis score (FPDS) and prevalence (FPDP), and Hockburn score (HBS) and prevalence (HBP). Significant differences between treatment groups (ANOVA, p <0.05). Different letters denote significant differences using DMRT grouping (p <0.05). SEM: Standard error of the mean, MSE: Error Mean Sum of Squares; F-stat (ANOVA F-statistic).

|  | T1 | T2 | T3 | T4 | T5 | T6 | SEM | MSE | F-stat | P-val |
| --- | --- | --- | --- | --- | --- | --- | --- | --- | --- | --- |
| C0 |  |  |  |  |  |  |  |  |  |  |
| BW | 2.435 | 2.405 | 2.436 | 2.397 | 2.383 | 2.376 | 0.011 | 0.005 | 0.819 | 0.546 |
| FCR | 1.368<sup>b</sup> | 1.393<sup>ab</sup> | 1.389<sup>ab</sup> | 1.418<sup>a</sup> | 1.410<sup>a</sup> | 1.412<sup>a</sup> | 0.005 | 0.001 | 2.891 | **0.030** |
| ADG | 60.876 | 60.118 | 60.902 | 59.933 | 59.580 | 59.410 | 0.282 | 2.949 | 0.819 | 0.546 |
| MT | 3.500 | 2.867 | 3.133 | 3.167 | 3.033 | 3.167 | 1.249 | 1.249 | 0.209 | 0.956 |
| EPEF | 406.007 | 398.163 | 402.015 | 389.479 | 390.510 | 388.676 | 2.662 | 244.196 | 1.312 | 0.285 |
| FPDS | 0.027 | 0.020 | 0.027 | 0.010 | 0.007 | 0.010 | 0.003 | 0.000 | 1.440 | 0.239 |
| FPDP | 2.667 | 2.000 | 2.667 | 1.000 | 0.667 | 1.000 | 0.314 | 3.333 | 1.440 | 0.239 |
| HBS | 0.010 | 0.013 | 0.007 | 0.000 | 0.007 | 0.003 | 0.002 | 0.000 | 1.200 | 0.333 |
| HBP | 1.000 | 1.333 | 0.667 | 0.000 | 0.667 | 0.333 | 0.178 | 1.111 | 1.200 | 0.333 |
| C1 |  |  |  |  |  |  |  |  |  |  |
| BW | 2.417 | 2.418 | 2.439 | 2.412 | 2.378 | 2.436 | 0.0101 | 0.004 | 0.731 | 0.606 |
| FCR | 1.425 | 1.426 | 1.415 | 1.447 | 1.436 | 1.426 | 0.004 | 0.001 | 1.146 | 0.358 |
| ADG | 58.945 | 58.967 | 59.480 | 58.832 | 58.009 | 59.419 | 0.248 | 2.300 | 0.731 | 0.606 |
| MT | 1.767 | 2.200 | 2.600 | 2.867 | 1.833 | 2.433 | 0.128 | 0.501 | 2.250 | 0.075 |
| EPEF | 383.464 | 383.409 | 386.183 | 371.883 | 377.150 | 381.341 | 2.150 | 167.042 | 0.975 | 0.449 |
| FPDS | 0.020 | 0.010 | 0.043 | 0.020 | 0.017 | 0.027 | 0.004 | 0.001 | 0.516 | 0.762 |
| FPDP | 1.667 | 1.000 | 3.333 | 2.000 | 1.333 | 2.000 | 0.308 | 3.556 | 0.725 | 0.610 |
| HBS | 0.030 | 0.027 | 0.033 | 0.037 | 0.023 | 0.023 | 0.005 | 0.001 | 0.973 | 0.450 |
| HBP | 2.000 | 2.000 | 2.667 | 2.667 | 2.000 | 1.667 | 0.312 | 3.533 | 0.962 | 0.456 |
| C2 |  |  |  |  |  |  |  |  |  |  |
| BW | 2.648 | 2.625 | 2.629 | 2.603 | 2.586 | 2.597 | 0.013 | 0.006 | 0.540 | 0.744 |
| FCR | 1.398<sup>d</sup> | 1.428<sup>bc</sup> | 1.417<sup>c</sup> | 1.459<sup>a</sup> | 1.442<sup>a</sup> | 1.442<sup>ab</sup> | 0.004 | 0.000 | 17.473 | **0.000** |
| ADG | 64.591 | 64.026 | 64.120 | 63.477 | 63.062 | 63.338 | 0.307 | 3.633 | 0.540 | 0.744 |
| MT | 2.000 | 2.567 | 2.900 | 2.033 | 2.800 | 2.367 | 1.011 | 0.166 | 0.857 | 0.521 |
| EPEF | 429.986<sup>a</sup> | 412.178<sup>bc</sup> | 416.671<sup>ab</sup> | 401.411<sup>c</sup> | 399.630<sup>c</sup> | 403.987<sup>bc</sup> | 2.557 | 141.778 | 5.617 | **0.001** |
| FPDS | 0.230 | 0.243 | 0.213 | 0.260 | 0.263 | 0.260 | 0.012 | 0.006 | 0.401 | 0.844 |
| FPDP | 17.333 | 17.667 | 16.667 | 19.333 | 19.333 | 19.000 | 0.945 | 36.222 | 0.218 | 0.952 |
| HBS | 0.153<sup>b</sup> | 0.153<sup>b</sup> | 0.200<sup>ab</sup> | 0.243<sup>a</sup> | 0.190<sup>ab</sup> | 0.210<sup>ab</sup> | 0.009 | 0.002 | 3.214 | **0.019** |
| HBP | 13.000<sup>b</sup> | 12.000<sup>b</sup> | 15.333<sup>ab</sup> | 17.667<sup>a</sup> | 14.000<sup>ab</sup> | 15.667<sup>ab</sup> | 0.562 | 9.133 | 2.727 | **0.038** |

**Table 2.**
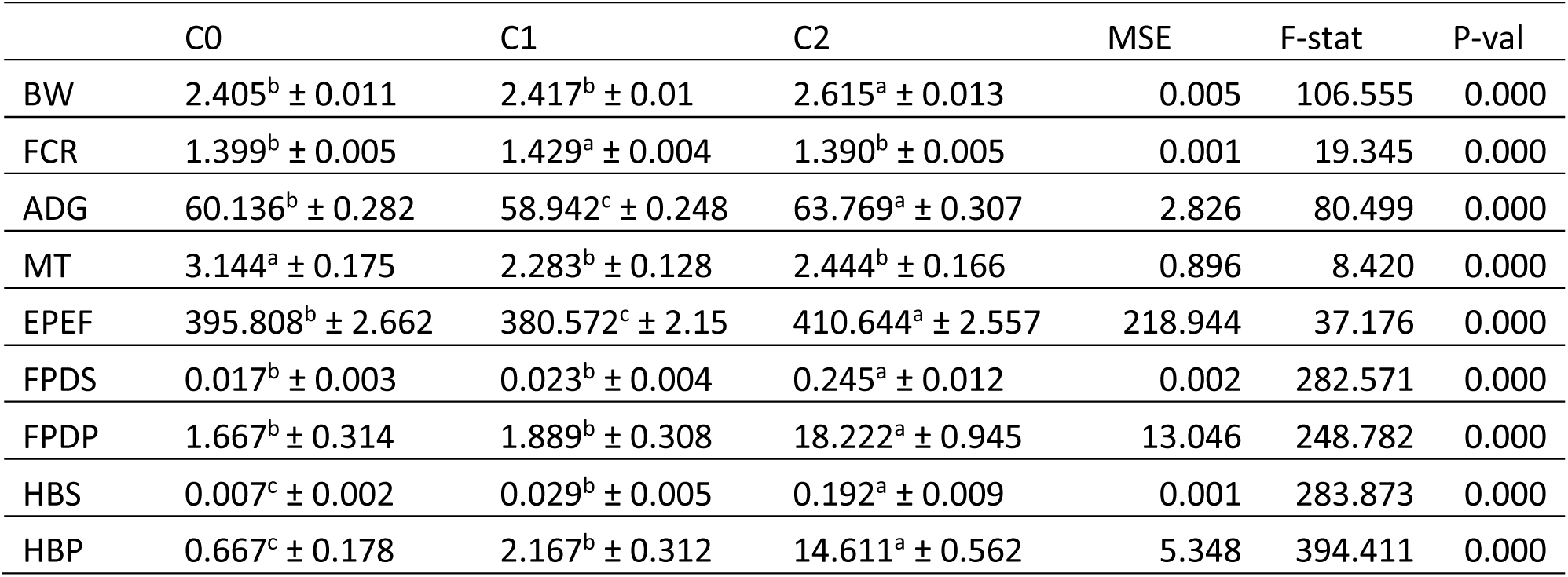
Performance and health parameters of birds across cycles. Cycle 0 (C0), Cycle 1 (C1) and Cycle 2 (C2) (N = 108 pens, 36 pens/cycle). Performance parameters taken as mean average per pen: Bird weight (BW), Average daily gain (ADG) - grams per day, Feed conversion ratio (FCR), Total mortality (MT), (E) European Production Efficiency Factor (EPEF), Footpad dermatitis score (FPDS) and prevalence (FPDP), and Hockburn score (HBS) and prevalence (HBP). Significant differences between treatment groups (ANOVA, p <0.05). Different letters denote significant differences using DMRT grouping (p <0.05). MSE: Error Mean Sum of Squares; F-stat (ANOVA F-statistic).

|  | C0 | C1 | C2 | MSE | F-stat | P-val |
| --- | --- | --- | --- | --- | --- | --- |
| BW | 2.405<sup>b</sup> ± 0.011 | 2.417<sup>b</sup> ± 0.01 | 2.615<sup>a</sup> ± 0.013 | 0.005 | 106.555 | 0.000 |
| FCR | 1.399<sup>b</sup> ± 0.005 | 1.429<sup>a</sup> ± 0.004 | 1.390<sup>b</sup> ± 0.005 | 0.001 | 19.345 | 0.000 |
| ADG | 60.136<sup>b</sup> ± 0.282 | 58.942<sup>c</sup> ± 0.248 | 63.769<sup>a</sup> ± 0.307 | 2.826 | 80.499 | 0.000 |
| MT | 3.144<sup>a</sup> ± 0.175 | 2.283<sup>b</sup> ± 0.128 | 2.444<sup>b</sup> ± 0.166 | 0.896 | 8.420 | 0.000 |
| EPEF | 395.808<sup>b</sup> ± 2.662 | 380.572<sup>c</sup> ± 2.15 | 410.644<sup>a</sup> ± 2.557 | 218.944 | 37.176 | 0.000 |
| FPDS | 0.017<sup>b</sup> ± 0.003 | 0.023<sup>b</sup> ± 0.004 | 0.245<sup>a</sup> ± 0.012 | 0.002 | 282.571 | 0.000 |
| FPDP | 1.667<sup>b</sup> ± 0.314 | 1.889<sup>b</sup> ± 0.308 | 18.222<sup>a</sup> ± 0.945 | 13.046 | 248.782 | 0.000 |
| HBS | 0.007<sup>c</sup> ± 0.002 | 0.029<sup>b</sup> ± 0.005 | 0.192<sup>a</sup> ± 0.009 | 0.001 | 283.873 | 0.000 |
| HBP | 0.667<sup>c</sup> ± 0.178 | 2.167<sup>b</sup> ± 0.312 | 14.611<sup>a</sup> ± 0.562 | 5.348 | 394.411 | 0.000 |

To evaluate the overall effects of the individual GHI components across the three cycles, we employed LASSO regression. As shown in Table 3, all GHI components were revealed to be significant predictors in at least one of the performance parameters. Notably, EPEF was largely influenced by all variables while MT was not associated with any. Analysis showed that ionophore administration at all stages can largely increase EPEF, but slightly decrease FCR, while the opposite is demonstrated by administration of ionophores at only during finisher stage. Vaccination was shown to have only slight negative effect on EPEF, while use of Probiotic A is shown to have negatively affect almost all parameters. In contrast, use of Probiotic B is generally associated with positive changes in respect to all parameters, apart from FCR and MT. The remaining predictors were shown to have large negative impacts on EPEF but very low positive impact on FCR.

**Table 3.** Lasso regression of performance and health parameters of broilers across all cycles (C0, C1 and C2), against sources of variability where the treatment components are coded as 1/0, corresponding to “Yes”/”No”. The model assumes an L1 penalty term which forces the beta coefficient of insignificant predictors to become zero. Significant beta coefficients in blue and red, indicate decrease and increase in performance parameter, respectively. Average daily gain (ADG) - grams per day, Bird weight (BW), European Production Efficiency Factor (EPEF), Feed conversion ratio (FCR), FPDP (Footpad dermatitis prevalence), HBP (Hockburn prevalence), Mortality (MT %)

| Predictors | Performance parameters |  |  |  |  |  |  |
| --- | --- | --- | --- | --- | --- | --- | --- |
|  | ADG | BW | EPEF | FCR | FPDP | HBP | MT |
| **(Intercept)** | **61.207** | **2.489** | **397.937** | **1.400** | **7.354** | **5.294** | **2.624** |
| Ionophore: All stages as Yes | 0.000 | 0.000 | 8.517 | -0.019 | 0.000 | 0.000 | 0.000 |
| Ionophore: Finisher only as Yes | 0.000 | 0.000 | -10.152 | 0.019 | 0.000 | 0.000 | 0.000 |
| Vaccinated: Yes | 0.000 | 0.000 | -0.002 | 0.000 | 0.000 | 0.000 | 0.000 |
| Probiotic A: Yes | -1.438 | -0.069 | -3.313 | 0.000 | -5.481 | -3.651 | 0.000 |
| Probiotic B: Yes | 1.931 | 0.099 | 17.376 | -0.018 | 10.534 | 9.644 | 0.000 |
| 1 % reduced crude protein: Yes | 0.000 | 0.000 | -13.893 | 0.034 | 0.000 | 0.000 | 0.000 |
| Essential oil: Yes | -0.427 | -0.017 | -10.748 | 0.022 | 0.000 | 0.000 | 0.000 |
| Prebiotic: Yes | 0.000 | 0.000 | -15.617 | 0.024 | 0.000 | 0.000 | 0.000 |

### Administration of different GHIs resulted in shifts in microbial community structure and diversity

The sequencing analysis of 118 caecal samples yielded a total of 2.6 B reads, from which a total of 84 MAGs with greater than 75% completeness and less than 5% contamination were recovered Specifically, 83 of these MAGs are detected in C1, while 78 are present in C2. All included MAGs were identified as bacterial species which represents 7 unique phyla, with Firmicutes_A as the most common designated phylum among all samples (Figures 2 and 3, Supplement file 2). This is followed by Firmicutes and Bacteroidota, while MAGs belonging to Proteobacteria was not present in any of the groups in C2 (Figure 2A). The global microbiota abundance is dominated by MAGs distributed across 57 identified genera accounting for 96% of the population. Among these, *Lactobacillus* is the predominant genus, while its member *Lactobacillus crispatus* is the most common species. However, predominant MAGs at genus and species levels were observed to largely vary across treatments and cycles (Figure 2, Supplement file 2). For instance, *Ruminococcus_G* was observed to have highest proportion in T1 and T3; *Anaerobutyricum* in T2, T5 and T6; and *Alisipes* for T6, while *Lactobacillus* was consistently highest in proportion among C2 treatment groups. Comparison of F/B ratio revealed significant differences between treatment groups in C1 only wherein T5 is observed to have a higher ratio than T4 (Figure 2D). Approximately 77% of the MAGs (n = 65) contained SCGs for phylogenetic mapping, showing 4 major groupings which are dominated by Firmicutes_A species. Among these MAGs, bin.108 (*CAG-267 sp001917135*) is revealed as the most novel species (Figure 4).

**Figure 2.**
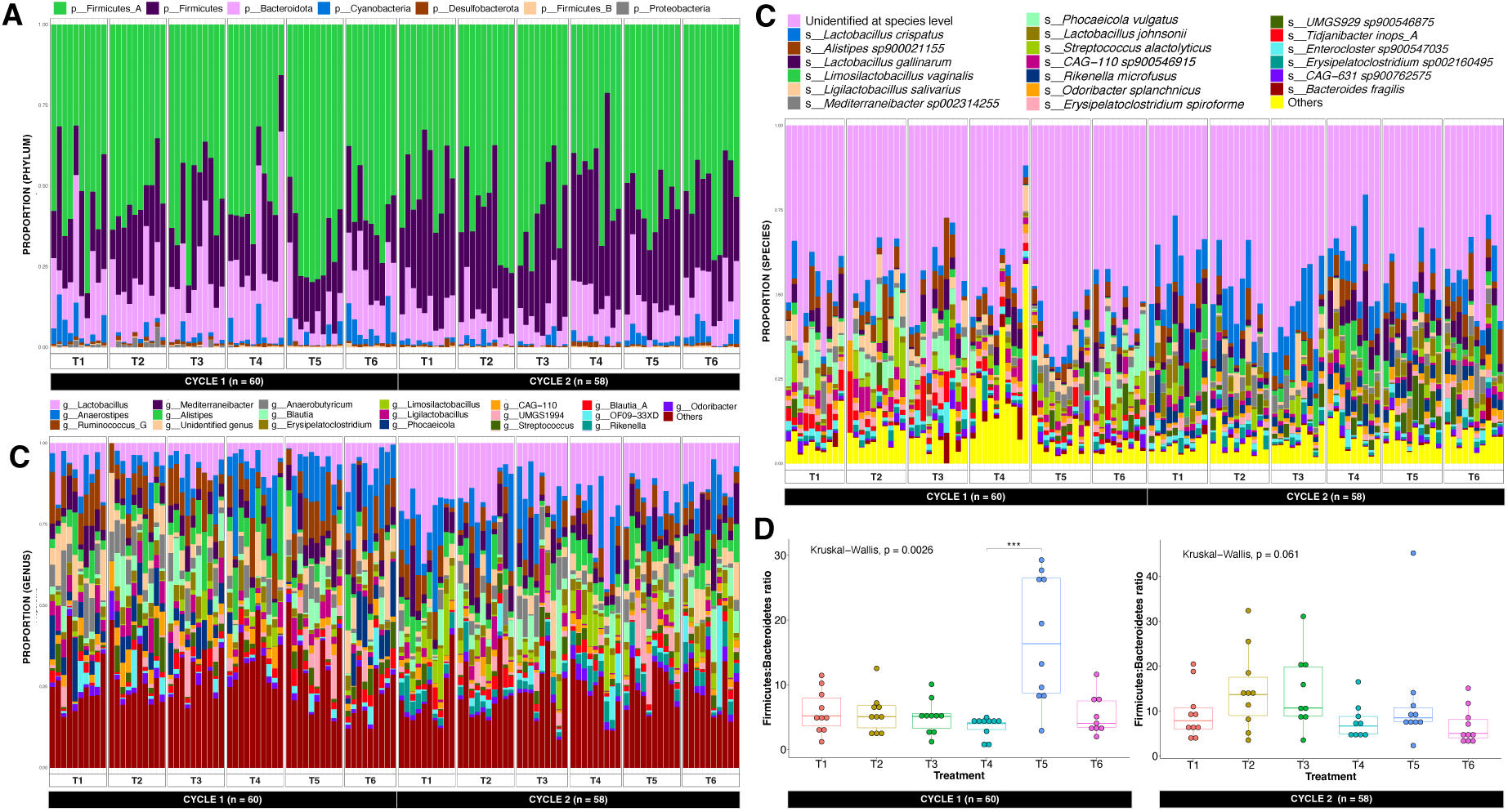
Sample-wise proportion of 84 MAGs recovered from C1 (n = 60) and C2 (n = 58). Proportions at (A) Phylum, (B) Genus (genera with <2% prevalence grouped into “Others”) and (C) Species levels (species with <1% prevalence grouped into “Others”), ranked from most dominant to least, grouped per treatment/cycle. Plot also shows (D) Firmicutes:Bacteroidota ratio, grouped per cycle and treatment, with significant differences based on Kruskal-Wallis (p < 0.05) and pairwise Dunn testing with Bonferroni correction: p < 0.001).

**Figure 3.**
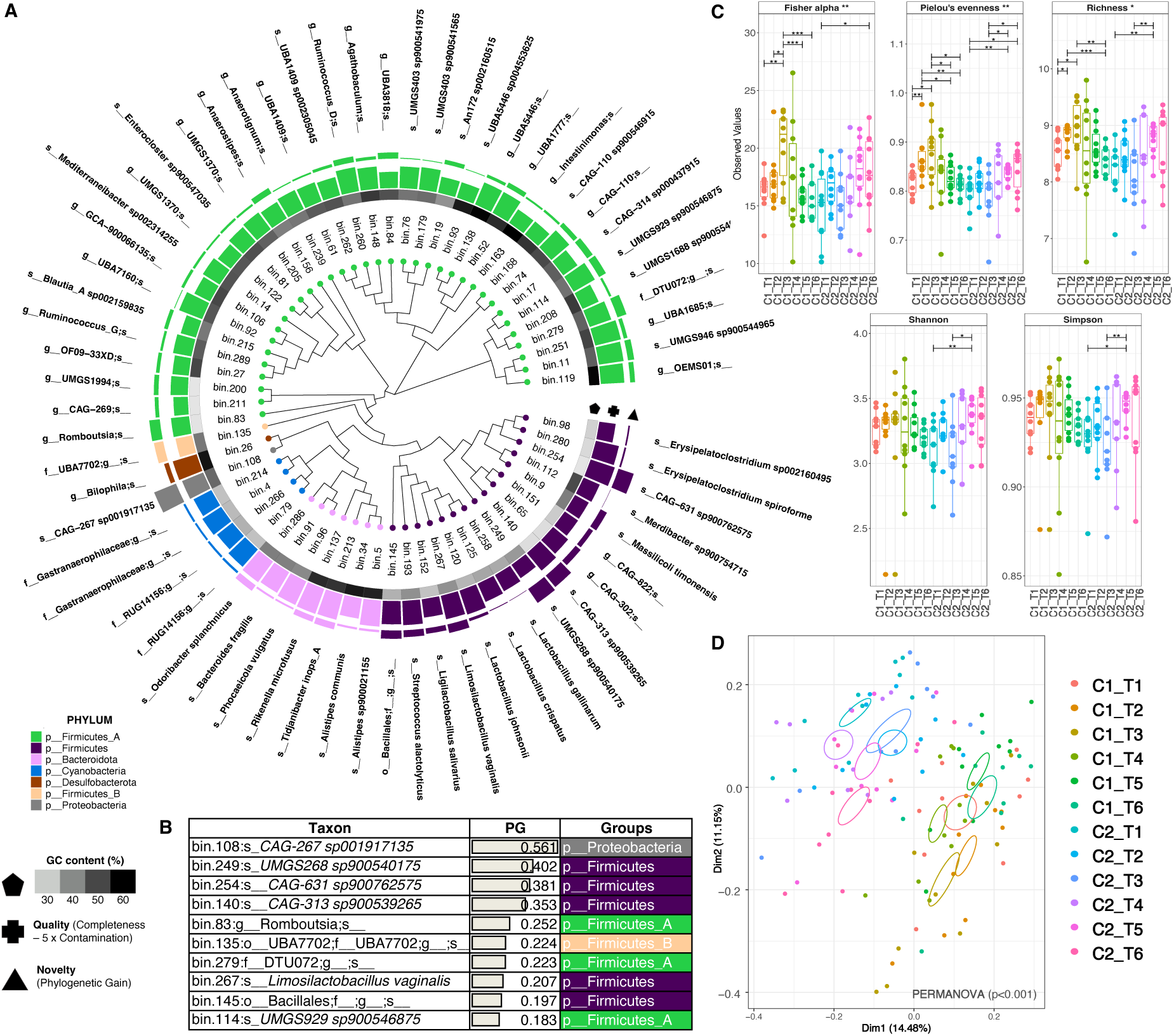
Phylogeny and taxonomic diversity of MAGs recovered from C1 (n = 60) and C2 (n = 58). (A) Phylogenetic tree of 65 MAGs recovered via GToTree using 25 bacterial and archaeal specific single copy genes. The tree also features G-C content, Quality index (genome completion – 5 x genome contamination), and Novelty (represented by phylogenetic gain (PG) values calculated using GTDB toolkit). (B) 10 most novel MAGs shown (indicated by high PG). Finally, alpha diversity (C) is represented by Fisher alpha, Pielou’s evenness, Rarefied richness, Shannon and Simpson index, with ANOVA significance: p < 0.001 (***), p < 0.01 (**), p < 0.05 (*), (D) Beta diversity is represented by PCoA plot of Bray-Curtis indices, with PERMANOVA (p <0.001).

**Figure 4.**
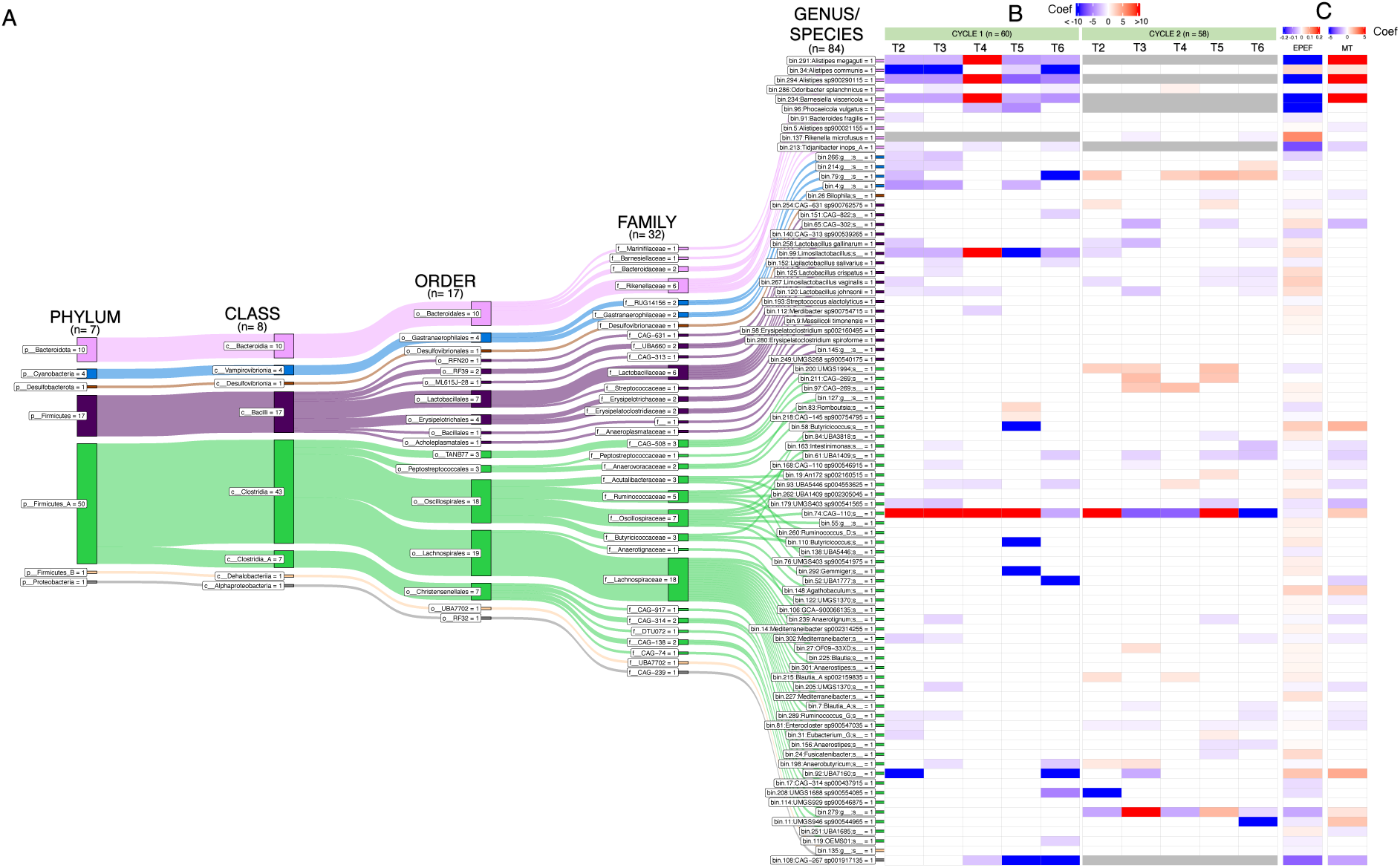
Taxonomic classification and differential analysis of 84 MAGs recovered from C1 (n = 60) and C2 (n = 58). (A) A sankey plot illustrating the classification of the bacterial species at various taxonomic ranks. The figure also includes general linear latent variable model (GLLVM) results showing association of MAGs with (B) treatment groups (with C1 and C2 done separately) and (C) performance and health parameters – EPEF (European Performance Efficiency Factor) and MT (Total mortality). Coef: GLLVM coefficients in blue (<0) or red (>0), representing negative or positive coefficients (or decrease or increase) for each MAG against GHI treatments (T2, T3, T4, T5, T6) as predictors, in comparison to T1 (reference). Coefficients in white indicates that coefficient is insignificant (no association), while grey indicates that the MAG was not recovered in that treatment.

Next, we hypothesised that GHI groups may influence in the microbial community diversity, which we then estimated through use of various alpha and beta diversity metrics (Figure 3). In C1, microbial communities of T3 samples were found richer and possessed more even distribution than other groups (T1, T2, T4, and T6) as indicated by significantly higher FA, R and PE values. Meanwhile in C2, the majority of alpha metrices of T5 and T6 were significantly elevated compared to T1 and T3. The principal components (PC1 and PC2) of our PCoA plot on Bray-Curtis estimates explained a considerable portion of the variance (14% and 11%, respectively), from which a discernible separation (PERMANOVA, p < 0.001, R^2^ = 27.84%) between C1 and C2 groups was observed. While separation between treatment groups is not distinct, slight clustering of groups can be observed: T2 and T3 (in C1) are separated from the other groups, while T1, T2 and T6 (in C2) are separated from the rest.

### Metagenomic-assembled genome composition is associated with administration of different GHIs and broiler performance

To further understand individual influence of GHI groups on microbiome and performance, we conducted GLLVM regression analysis of each MAG (using T1 as reference predictor). A total of 43 and 38 MAGs (59 for both) were observed to have a significant association with GHI groups in C1 and C2, respectively (Figure 4, Supplement file 1). For C1, the majority of these MAGs (n = 41/43, 95%) exhibited a negative association with GHI administration, denoting decrease in abundance compared to T1. From this, T2 had the greatest number of decreased MAGs, followed by T3 and T6, while increase of 5 MAGs were seen in T4. Notable MAGs to consistently change across groups in C1 were those belonging to *Bacteroidales* (bin.291 *Alistipes megaguti*, bin.34 *Alistipes communis*, bin.294 *Alistipes sp900290115*, bin.234 *Barnesiella viscericola*), as well as bin.99 (*Limolactobacillus*) and bin.74 (CAG-110). Bin.34 is also observed to have the lowest GLLVM coefficient, followed by bin.92 (g_*UBA7160*) and bin.4 (f_*Gastrophilaceae*). In addition, all Cyanobacteria MAGs were observed to show negative association with GHI groups.

In contrast, a higher number of positively associated MAGs (n = 15) was observed in C2 than in C1 (n = 7), from which T5 has the highest count (n = 8). Meanwhile, 25 MAGs were seen to have negative association with GHI groups in C2, with T6 having the highest number of negatively associated MAGs (n = 15). Bin.74 (*g_CAG-110*) is observed to be dynamic across all GHI groups, as well as the having the highest coefficient among all MAGs in C2; while bin.208 (*UMGS1688 sp90054085*) is shown to have the strongest negative association (Figure 4B).

For performance, we considered EPEF and MT to represent overall broiler health and performance (EPEF are highly correlated with the other measured performance parameters). Our analysis revealed that EPEF is positively associated with 40 MAGs and negatively associated with 33 MAGs, while MT is associated with only 30 MAGs (10 positive and 20 negative associated MAGs). Bin.137 *(Rikenella microfusus*) is noted to have the highest GLLVM coefficient against EPEF, followed by bin.92 *(g_UBA7160*) and bin.58 (*Butyricicoccus*). Bin.234 (*Barnesiella* viscericola) is observed to have the strongest negative association with EPEF but also the strongest positive association with MT (Figure 4C).

Interestingly, at least one of the *Lactobacillaceae* MAGs (bin.258 *Lactobacillus gallinarum*, bin.99 *Limolactobacillus*, bin.152 *Ligilactobacillus*, bin.125 *Lactobacillus crispatus*, bin.267 *Limolactobacillus salivarius*, bin.120 *Lactobacillus johnsonii*) are negatively associated across the GHI groups in both cycles (except for T4) but are also seen to have positive association with EPEF, indicating that decreased levels of *Lactobacillaceae* MAGs in GHIs may contribute to diminished EPEF values. Similar patterns were also seen for other MAGs such as bin.34, bin.137, bin.65, and bin.84. Furthermore, numerous MAGs enriched in GHI groups but negatively associated with EPEF were also identified (Cyanobacteria MAGs, bin.200: UMGS1993, bin.279: f_DTU072 in C2) which signifies that elevated levels of these MAGs can contribute to decreased levels of EPEF.

### Administration of different GHIs resulted in shifts in metabolic functions

Since significant differences in MAGs were observed across treatments, we then investigated the impact of GHIs on metabolic functions. Enzymes including CAZymes and proteases are important for metabolism and reproduction of microbial species, hence they may also play crucial roles in nutrition and digestive physiology of chickens. As seen in Figure 4, we detected a total of 128 CAZymes belonging to two major families – glycoside hydrolase (GH) and polysaccharide lyase (PL), with the former being more dominant. Both CAZyme families were significantly different in abundance across C1 treatment groups, revealing T5 to have highest GH abundance but also lowest PL abundance (p < 0.05, ANOVA) (Figure 5A). Meanwhile, there were no significant differences between groups within C2, but they have a relatively higher overall CAZyme abundance than C1 groups. Furthermore, clustering analysis exhibited separation of T4 and T5 from the other groups in C2, and T5 in C2 (Figure 5B).

**Figure 5.**
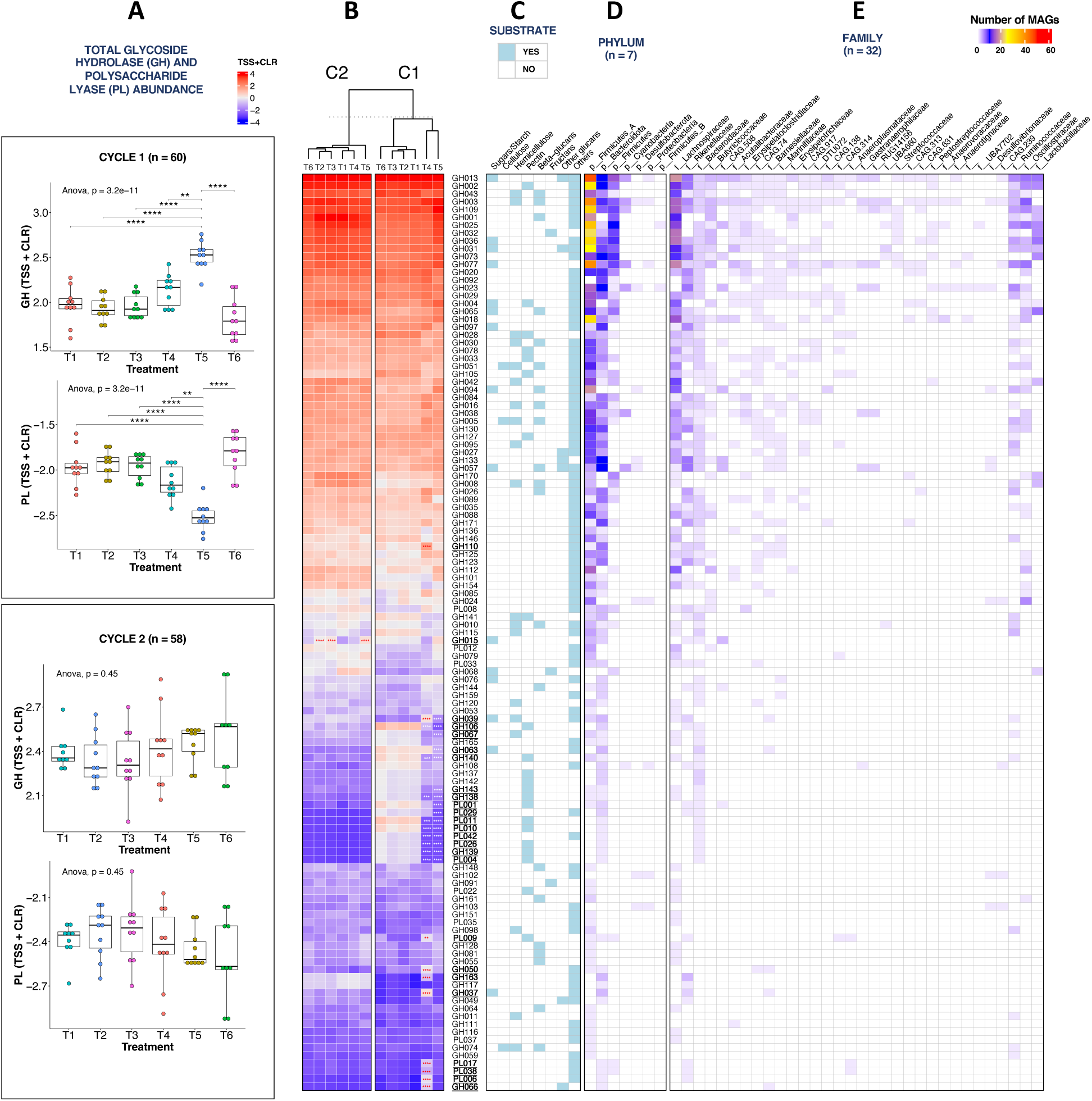
Carbohydrate-active enzymes (CAZyme) gene abundance recovered from C1 (n = 60) and C2 (n = 58). (A) Mean normalised abundance of CAZyme gene IDs; Red and blue colour of heatmap cells indicates high and low abundance, respectively); IDs in bold and underlined indicates significant increase or decrease in abundance compared to T1 based on Wald-Test *(DESEq2)*. CAZyme IDs grouped according to (B) substrate/function based on the dbCAN2 and MetaCyc databases) across treatment groups and cycle. Number of MAGs containing CAZyme genes, grouped according to (C) phylum and (D) family taxonomic ranks, (E) Comparison of total glycoside hydrolase (GH) and polysaccharide lyase (PL) abundances across treatment groups and cycles, based on ANOVA and *posthoc* pairwise t-testing with Bonferroni correction. Normalisation method: Total Sum Scaling and Centralised Log Ratio (TSS + CLR). Significance: p<0.0001 (****), p < 0.001 (***), p < 0.01 (**), p < 0.05 (*). White and red significance indicates decrease and increase of MAGs in treatments T2 to T6 compared to T1, respectively.

The most common enzyme across all treatments is observed to be GH013 which is detected in 27 bacterial families in our study. A total of 24 enzymes are noted to have significantly different abundance between T1 and GHI groups T4 and T5 (C1) (Wald Test, p <0.05, Figure 5B). For T4, 9 enzymes with activity against pectin were significantly lower than T1, while 11 enzymes with activity against starch/sugars and other carbohydrates are significantly higher. Similarly, 15 enzymes with activity against pectin and hemicellulose are downregulated in T5; these same enzymes are observed to be consistently present in the family Bacteroidaceae (Figure 5E). In C2, only GH15 is observed to have a significantly difference (upregulated in T2, T3, and T5) which has capacity for sugars/starch digestion.

Differential abundance of proteases across treatments was also investigated in this study. A total of 108 protease families distributed across 9 protease types were detected in our samples (Figure 6). Metallo peptidases were detected as the most common and most diverse protease catalytic type, with M38 family being the most dominant across 47 metallo families. Other detected catalytic types include inhibitors and threonine peptidases which were both significantly different in abundance in C1, where threonine abundance was specifically observed significantly lower in T4 compared to T1 (Dunn Test with Bonferroni correction, p <0.05). In C2, abundances of cysteine peptidases were markedly disparate (Kruskal Sum Rank test, p < 0.05), with T4 having the highest value among all groups but although insignificant. At family level, divergent clustering of T4 and T5 from other groups were seen in C1; In contrast, for C2, T1 and T4 are seen to be more similar in abundance than the other groups. Across all treatment groups, C38 (Cysteine) is consistently the most abundant peptidase, followed by M38 and S33. However, diminished levels of M28X were evident in T4 and T5 while elevated levels of M93 and M28A were observed in T4 in C1. Notably, all these differentiated families are present in Bacteroidaceae family (Figure 6E).

**Figure 6.**
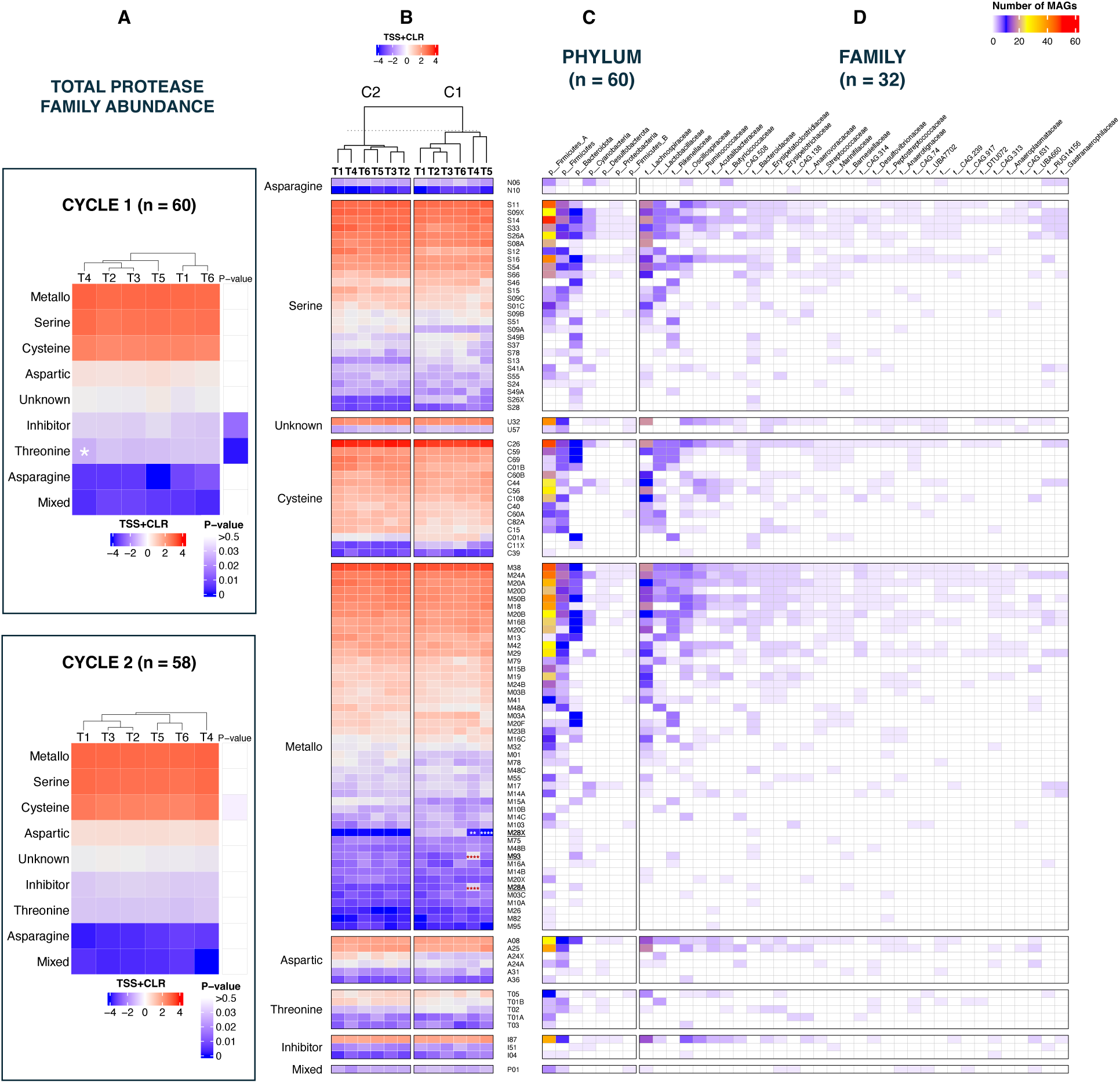
Mean normalised abundance of proteases across treatment groups and cycles. (A) Mean abundance of individual protease families (MEROPS ID) grouped according to protease type in accordance to MEROPS database [44]. MEROPS protease ID in bold and underlined indicates significant increase or decrease in abundance compared to T1, based on Wald-Test (using *DESEq2*). Number of MAGs with protease gene grouped according to taxonomic ranks (B) phylum and (C) family, (D) Mean abundance of total protease grouped according to family across groups and cycles, statistical differences were based on Kruskal-Sum Rank and *posthoc* pairwise Dunn testing with Bonferroni correction. Normalisation method: Total Sum Scaling and Centralised Log Ratio (TSS + CLR). Significance: p<0.0001 (****), p < 0.001 (***), p < 0.01 (**), p < 0.05 (*), white and red significance indicates decrease and increase of MAGs in GHI treatments (T2, T3, T4, T5, T6) compared to T1, respectively.

**Figure 7.**
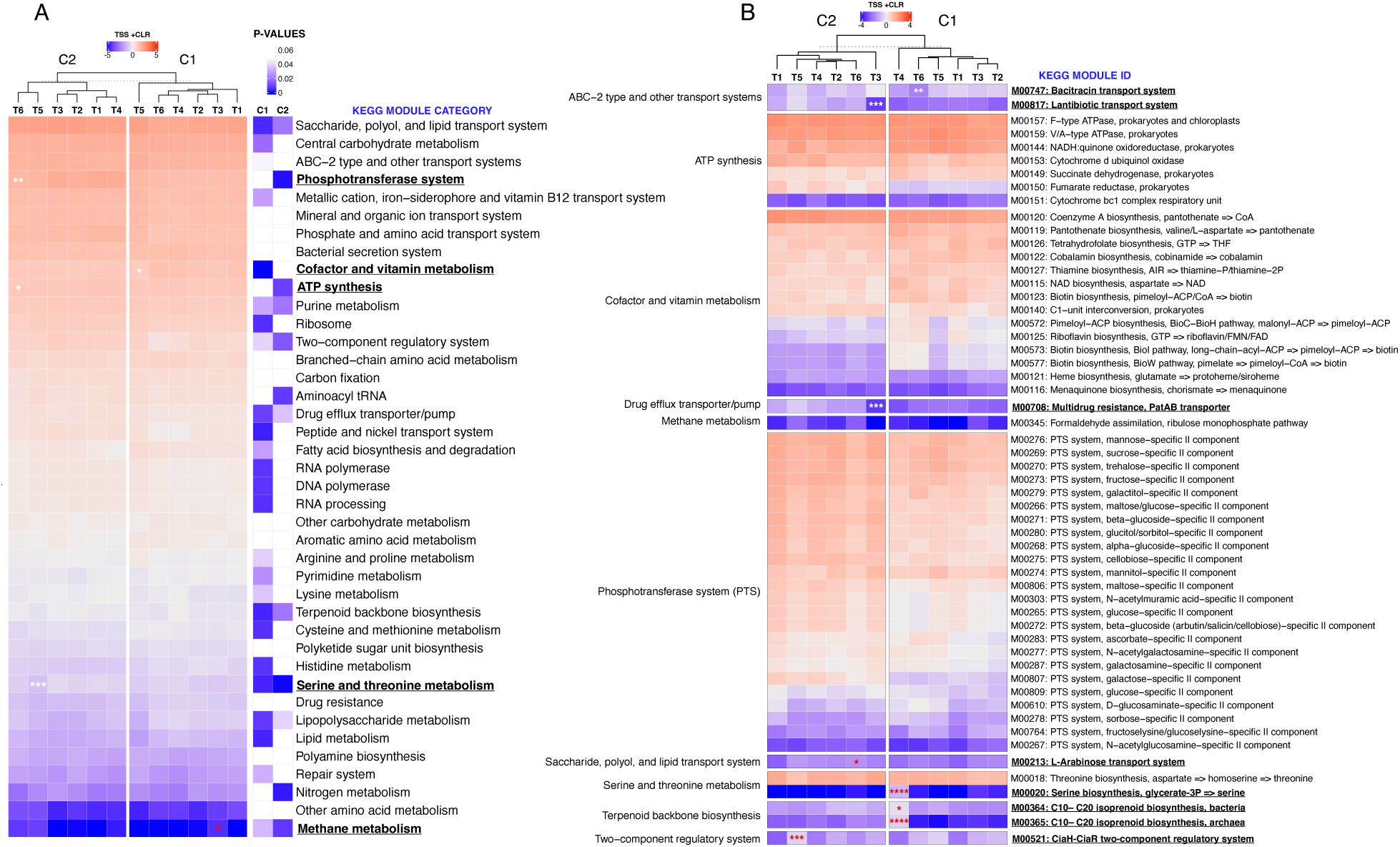
KEGG metabolic module abundances across treatment and cycles. Mean normalised abundance of (A) KEGG module categories and respective p-values based on Kruskal-Wallis testing (B) Selected KEGG modules across treatment groups and cycles (C1 and C2). Red and blue colour of heatmap cells indicates high and low abundance, respectively. KEGG category and module IDs in bold and underlined indicates significant difference in abundance compared to T1 based on *posthoc* pairwise Dunn testing with Bonferroni correction and DESEq2, respectively. Normalisation method: Total Sum Scaling and Centralised Log Ratio (TSS + CLR). Significance: p<0.0001 (****), p < 0.001 (***), p < 0.01 (**), p < 0.05 (*). Significance in white and red indicates decrease and increase of MAGs in GHI treatments (T2, T3, T4, T5, T6) compared to T1, respectively.

Gut microorganisms engage in complex metabolic interactions, potentially involving various production of metabolites that modulate host physiology. Based on our analysis, we determined that GHI administration has also an impact on other KEGG modules including various transport and transporter systems, amino acid metabolism and nucleic acid metabolism pathways (Figure 8).

**Figure 8.**
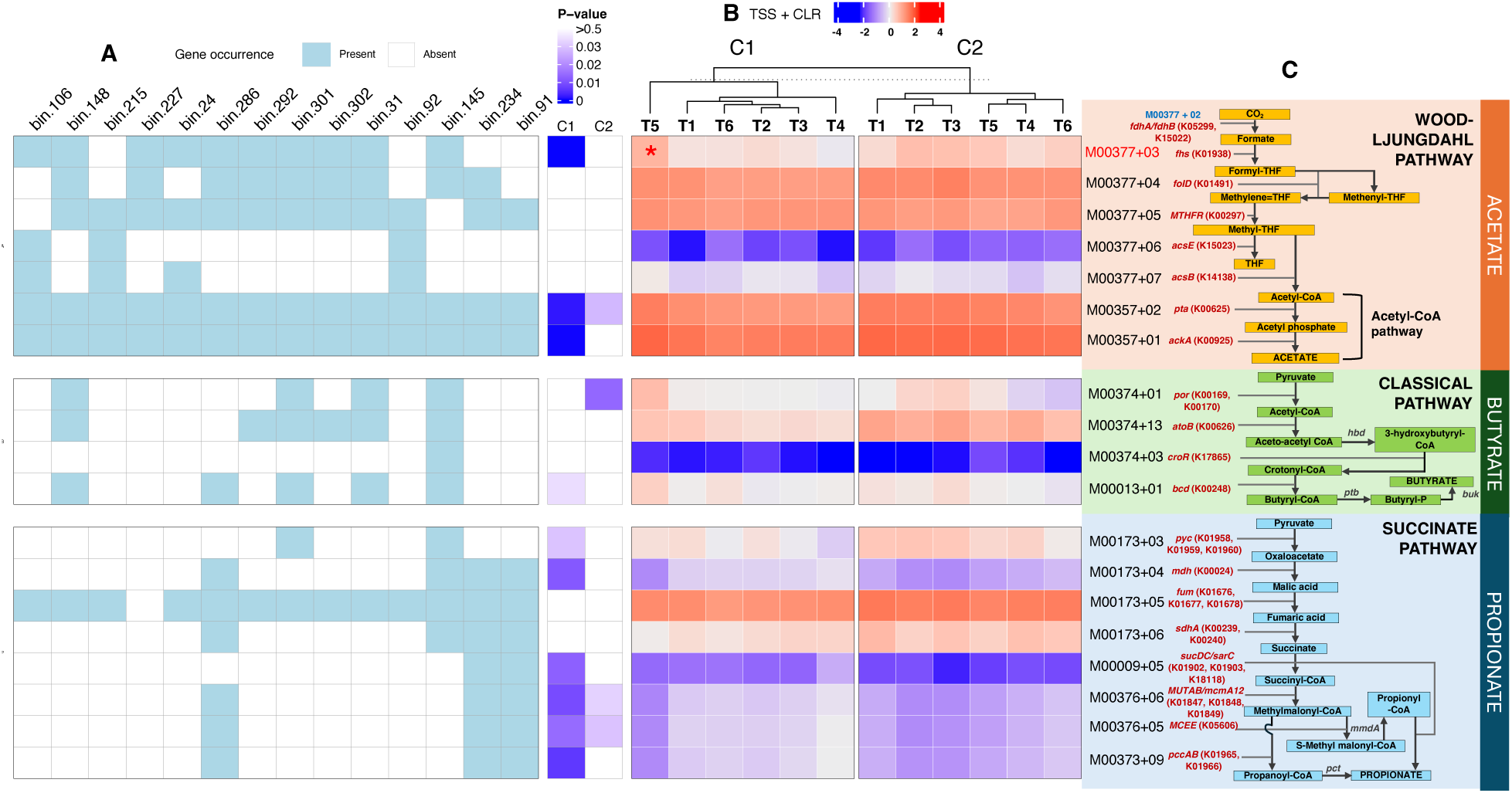
Metagenomic abundance of elements of short-chain fatty acids (SCFA) pathways. (A) Selected MAGs containing most complete SCFA submodules. Blue and white colour indicates presence and absence of element, respectively; (B) Heatmap of normalised abundances of SCFA enzyme corresponding module IDs across treatment groups and cycles (C1 and C2), with significance based on Kruskal-Rank Sum test (p <0.05 in blue). TSS + CLR: Total Sum Scaling and Centralised Log Ratio. Red and blue colour of heatmap cells indicates high and low abundance, respectively; (C) SCFA pathways showing KEGG orthology (KO) numbers and enzyme names (in red). Submodule and significance in red: significant increase based on pairwise Dunn test with Bonferroni correction (p < 0.05). Submodules in blue: not detected in metagenome dataset in this study. Enzyme names in black: no matching submodule ID in KEGG database [41]. Enzymes are listed as follows: *fdhA*: Formate dehydrogenase alpha subunit; *fdhB*: formate dehydrogenase beta subunit; *fhs*: formate tetrahydrofolate ligase; *foID*: methylenetetrahydrofolate dehydrogenase; *MTHFR*: methylenetetrahydrofolate reductase; *acsE*: 5-methyltetrahydrofolate corrinoid/iron sulfur protein methyltransferase; *acsB*: acetyl-CoA synthase; *pta*: phosphate acetyltransferase; *ackA*: acetate kinase; *por*: pyruvate ferredoxin oxidoreductase; *atoB*: acetyl-CoA C-acetyltransferase; *hbd*: 3-hydroxybutyryl-CoA dehydrogenase; *croR*: 3-hydroxybutyryl-CoA dehydratase; *bcd*: butyryl-CoA dehydrogenase; *ptb*: phosphate butyryltransferase; *pyc*: pyruvate carboxylase; *buk*: butyrate kinase; *mdh*; malate dehydrogenase; *fum*: fumarate hydratase; *sdhA*: succinate dehydrogenase; *sucD*: succinyl-CoA synthetase alpha subunit; *sucD*: succinyl-CoA synthetase beta subunit; *aarC*: succinyl-CoA:acetate CoA-transferase; *MUTAB*; methylmalonyl-CoA mutase alpha and beta; *mcmA1*: methylmalonyl-CoA mutase, N-terminal domain; *mcmA2*: methylmalonyl-CoA mutase, C-terminal domain; *mmdA*: methylmalonyl-CoA decarboxylase subunit alpha; *MCEE*: epi; methylmalonyl-CoA/ethylmalonyl-CoA epimerase; *pccA*: propionyl-CoA carboxylase alpha chain; *pccB*: propionyl-CoA carboxylase beta chain; *pct*: propionate CoA-transferase.

Significantly varied abundances across treatment groups in 25 and 15 different KEGG function module categories in C1 and C2 were observed (p < 0.05, Kruskal-Rank Sum test). Heatmap clustering also exhibits the divergence of T6 from the other groups in C1, whilst both T5 and T6 have partitioned from the others in C2. However, compared to T1, abundance in GHI groups were significantly downregulated in 4 module categories namely phosphotransferase system and ATP synthesis in C2 – T6, serine and threonine metabolism in C2 – T5, and co-factor and vitamin metabolism in C1 – T5, while methane metabolism function is upregulated in C1 – T3. Furthermore, differential analysis (T1 as reference) of individual modules revealed decreased abundance of antibiotic related transport systems in GHI groups C1 – T6 (M00747) and C2 – T3 (M00817, M00708) and increase of organic compound biosynthesis and transport, and regulatory system modules in C1 – T4 (M00020, M00364, M00365), C2 – T6 (M00213), C2 – T5 (M00521).

### Administration of different GHI can impact short-chain fatty acid production pathways

Due to the known relevance of metabolic pathways involved in the production of short-fatty acids (SCFA) including acetate, butyrate and propionate on gut health, we also explored the effects of GHI on SCFA production. As shown in Figure 9, the majority of KEGG submodules related to SCFA production were present in our samples (excluding M00377+02). We also identified MAGs to have the complete sets of modules for Acetyl-CoA pathway (48/84 MAGs) and butyrate pathways (bin.145); However, none of our MAGs possessed complete sets of submodules for Wood-Ljungdahl and propionate pathways. Bin.92 and bin.215 have the highest number of submodules (n = 5), while MAGs bin.91 and bin.234 are missing M00173+03 – submodule for pyruvate carboxylase (*pyc*).

**Figure 9.**
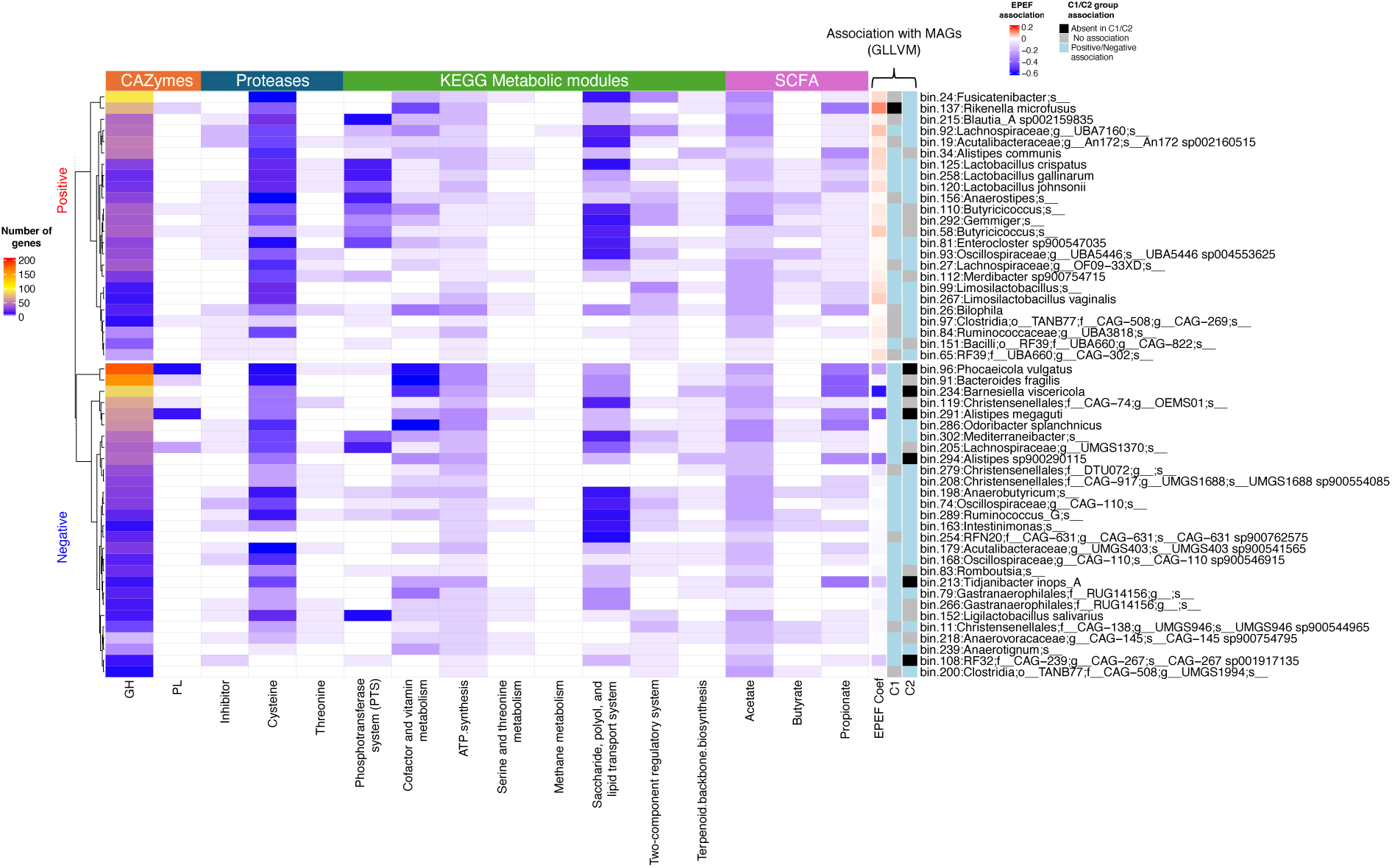
Overview of significant metagenomic features of 52 MAGs associated with EPEF and GHIs. EPEF: European Poultry Efficiency Factor, GH: Glycoside hydrolase, GHI: Gut health interventions, GLLVM: Generalised latent linear variable model, MAG: Metagenome-assembled genomes, SCFA: Short chain fatty acids, PL: Polysaccharide lyase.

Across our treatment groups, 11 out of the included 19 submodules are significantly varied, wherein propionate submodules were observed as mostly affected. From this, *post hoc* pairwise comparison revealed a single elevation of M00377+03 in C1-T5. Furthermore, distinct separation of T5 in C1 and distancing of T1 and T6 from other groups in C2 were observed.

### MAGs associated with better performance have higher capacity for nutrient digestion and metabolism

Based on above analysis, we explored the gene abundance of several metagenomic features (which were selected based on statistical significance among treatment groups) of 52 MAGs that were revealed to be associated with both EPEF and composition in GHI groups (Figure 9). Clustering analysis showed that MAGs with positive association with EPEF (EPEF+) have an overall higher number of genes encoding for the various significant metabolic features. Specifically, the EPEF+ group demonstrated higher genes in the PTS module, Butyrate and Propionate SCFA module while similar abundance in CAZymes and proteases between the two EPEF groups can be seen. However, it is interesting to point out that *Rikenella microfusus,* the MAG with highest GLLVM coefficient for EPEF, do not possess any genes for PTS and butyrate production modules.

## Discussion

The microbial community structure of the gut has been proven crucial in both host health and performance [1,8]. As such, supplementation of feed additives to diets and administration of other health interventions have become common approaches to modulate intestinal microbial communities and ensure the ideal growth and health of broilers [66]. Previously, we reported alterations in the chicken gut microbiome in response to various production systems, using 16S rRNA sequencing. Our findings revealed that inclusion of omega-3 in feed can result in increase in bacterial genera associated with short-chain fatty acid (SCFA) production and affect levels of pathogenic bacteria including *Campylobacter* levels through competitive exclusion [9]. Due to increasing demand for poultry meat worldwide [67], there is a need to optimise gut health for improved feed efficiency and overall health of broilers. Therefore, we designed modified gut health schemes across three broiler production cycles and assessed their influence on the caecal microbiome using metagenomic shotgun sequencing. Herein, we employed assembly of 84 high quality MAGs recovered from 118 caecal samples and analysis of metabolic function capacity of the MAGs and association with performance parameters and health.

Administration of ionophores and GHIs employed in this study have been previously reported to modulate gut microbiota, with documented growth-promoting effects and microbial alterations dependent on the specific formulation and application level (Cai et al., 2022; Maki et al., 2019; Robinson et al., 2019; B. Wang et al., 2021). Within our study, we observed variations across the three cycles, where C2 groups demonstrated best performance. Since Probiotic B was revealed to have the highest positive impact on EPEF, we posit that the efficacy of Probiotic B may be a key contributing factor to the comparatively superior performance observed in cycle 2 compared to the other cycles provided with Probiotic A. These findings further underscore the importance of strain-specific effects and dosage considerations when implementing probiotic interventions in poultry systems, as reported before [68,69]. However, cycle variations within the same production system have also been reported previously and are hypothesised to occur due to variations in climate, management, and fluctuations in microbiota of day-old chicks as affected by hatchery-to-farm transfer [70]. Furthermore, some differences in MAG composition between C1 and C2 brought upon by differences in the GHIs used may explain their disparities in performance. For instance, *Rikenella microfusus* (bin.137), positively associated with EPEF but absent in C1, has been identified for its potential probiotic effects, attributed to its role in producing short-chain carboxylic acids, which contribute to maintaining cell structure integrity [71]. Conversely, *Alistipes spp.* (bin.291, bin.294) and *Phocaeicola vulgatus* (bin.96), all observed to be negatively associated with EPEF and present in only C1, have also been implicated in human health issues such as cancer, cardiovascular disorders and inflammatory related diseases [72,73]. Another noteworthy MAG is *Barnesiella viscericola* (bin.234), previously reported as an efficient coloniser of chicken caeca [74], but here observed as negatively correlated with EPEF but positively correlated with MT, and absent in C2.

Comparison of groups have also revealed substantial differences in microbial diversity and composition between GHI groups and the control (T1). Specifically, our analysis revealed that T1 and T3 to have better overall performance but exhibited lower alpha diversity than other groups (in C2). According to Coyte et al., (2015), high alpha diversity in the gut microbiome tends to destabilize microbiome communities, potentially leading to decreased ecological stability which is the ability to return to a natural state after a perturbation. Unstable gut microbial communities are then less likely to maintain beneficial symbiotic relationships and may be more susceptible to disturbances or shifts that could impact the host’s health and productivity [75]. With this, the moderate microbial diversity shown in T1 and T3 may indicate a more balanced and stable gut, thereby becoming supportive of an optimal performance. Furthermore, these discrepancies can be explained by MAGs with association with performance that were differentiated among groups. MAGs belonging to Lactobacillaceae (bin.258 *L. gallinarum,* bin.99 *Limosilactobacillus,* bin.125 *L. crispatus*, bin.120 *L. johnsoniiI*), Butyricicoccaceae (bin.58, bin.110 *Butyricioccus spp.)*, Ruminococcaceae (bin.292 *Gemmiger*) and Lachnospiraceae (bin.81 *Enterocloster sp900547035*, bin.24 *Fusicatenibacter*), which are families known for their beneficial roles in gut health such as SCFA production, gut integrity promotion, and protection against pathogens such as *Salmonella* spp. [76–81], were identified beneficial for performance but were decreased in different GHI groups in comparison to T1. Meanwhile, MAGs including bin.254 CAG-631 sp900762575, bin.200 UMGS1994, bin.83 *Romboutsia*, and bin.74 CAG-110, were enriched in GHI groups but are negatively associated with EPEF. Similarly, increase in *Romboutsia* has also been noted in broilers given *Bacillus subtilis* and a coccidiosis vaccine [82] and reported to also negatively impact performance of breeder broilers [83].

The standard diet of broilers usually consists of approximately 70% carbohydrates, encompassing starch, oligosaccharides, and non-starch polysaccharides (NSP) like cellulose, hemicellulose as well as pectin [84]. These NSPs remain undigested by the host, serving as substrates for the gut microbiome. Consequently, gut microorganisms possess a diverse range of genes encoding enzymes known as CAZymes, which facilitate the breakdown and metabolism of these polysaccharides [85]. CAZymes are categorized into families, including Polysaccharide lyases (PL) and glycoside hydrolases (GH), based on sequence similarities, although members within the same family may exhibit different substrate specificities [85]. In our study, we observed preference of gut microbiota in T4 for “other glucans” and “other glycans” and depletion of pectin bacterial specialists in T4 and T5 of C1. Since availability of readily accessible growth substrates diminishes as it passes through the gastrointestinal tract [86], we hypothesise that due to the limited availability of protein (amino acids) in T4, more preferred substrates such as starch and pectin has been digested in the upper intestines, leaving caecal microbiota to use other glycans as substrate. Meanwhile, depletion in T5 may be explained by the reduction of *Bacteroidia* bacteria (in T5), similarly reported by Ding et al., (2022), which are microorganisms shown here to digest pectin.

Digestion of protein available in the diet is also of great importance for optimisation of gut health [88]; However, there is limited information on its association with gut microbial functions in poultry. Previous research has shown that approximately 20% of crude protein (CP) taken in by broilers goes undigested due to insufficient concentrations of endogenous proteinases in the host [86,89]. Consequently, undigested protein (or ileal bypass protein) which are fermented by gut microbiota in the hindgut (caeca), can encourage with increased growth of *Clostridium perfringens* and production of detrimental metabolites including ammonia, indoles, and phenols [86]. Hence, we included reduction of CP as one of our gut health approaches in our study (as represented by T4). Nonetheless, this group was shown to have significantly lower overall performance. Since restriction of CP might have resulted in decrease of threonine [90,91], observed deficiency of threonine proteases in T4 may have affected threonine intestinal absorption by the host. A large proportion of host dietary threonine, known as the second (or third) limiting amino acid in broilers, is predominantly used by the host for production of mucin, an important glycoprotein that preserves the integrity of intestinal mucosa and function (Qaisrani et al., 2018). With this, it is hypothesised that there could be impaired intestinal permeability in T4 broilers which may have then contributed to overall poorer nutrient absorption, thereby affecting growth and performance. In addition, T4 in C1 has also been revealed to have differentially abundant genes for metalloproteases, a family of peptidases previously linked to have overactivity in patients with irritable bowel syndrome (Mills et al., 2022). Meanwhile, the elevated levels of HB metrics in T4 of C2 was unexpected given that reduction of CP in diets has commonly been associated with lower incidence of footpad lesions and better litter quality [94,95]. However, significances in abundance among C2 groups were determined for cysteine proteases, which are proteases renowned for their involvement in virulence and their ability to induce inflammatory responses including atopic dermatitis in humans [96,97]. It is interesting to note that C2 groups have both higher cysteine and HB levels than C1 groups (especially C01A), also indicating possible link between cysteine levels and HB occurrence.

The gut microorganisms participate in the metabolism and uptake of numerous nutrients and play crucial roles in preserving the integrity of the intestinal barrier, regulating the immune system, and protecting against pathogen colonisation [2]. In this study, we primarily identified differentiation of metabolic functions involved in energy production, nucleotide metabolism and drug transport related pathways among treatment groups. This is in line with previous research that have shown the following: 1) Probiotic supplementation in broilers can affect vitamin biosynthesis and other energy related metabolic activities [98]; 2) Caecal microbial changes due to probiotics can affect emissions such as nitrogen or ammonia [99]; 3) Antimicrobials can disrupt the nucleotide pool of bacterial cells, resulting in increased nucleotide biosynthesis and elevated central carbon metabolism [100]; and 4) By influencing the expression of microbial enzymes involved in pathways linked to nucleotide, amino acid, carbohydrate, and energy metabolism, antimicrobials could potentially steer metabolic flow, regulating bacterial proliferation or generating metabolites that affect the host [101]. Specifically, we observed reduction of the phosphotransferase system, cofactor and vitamin metabolism, and serine and threonine modules, in several GHI groups (T5 and T6) compared to T1, which are pathways involved in nutrient absorption and defence against infection in the host [102–104]. Increase in methane metabolism was also detected in T3 of C1 which was also similarly reported in other studies involving probiotic use in chickens [105,106]. However, this is only attributed to one KEGG module (M00345) which was detected in bin.92 UBA7160 (Lachnospiraceae). We also observed the relative decrease in drug transport-related modules namely bacitracin, lantibiotic and PatAB transport systems. Further research is needed to confirm whether this can be explained by the similarity in pharmacological mechanisms of ionophores to bacitracin and lantibiotics, both of which are antimicrobials that also prevent cell wall synthesis and mainly act on gram-positive bacteria [107,108].

SCFA, metabolites synthesised by caecal gut microorganisms from breakdown of dietary fibre, play vital roles in improving metabolism, facilitating nutrient digestion and absorption, thereby promoting optimal health, growth, and well-being in poultry [2]. From our analysis, we generally observed gene differentiation of SCFA modules among our treatments, wherein T5 was observed to have relatively higher gene abundance among other groups, especially in acetate production. We speculate this can be due to the effect of essential oils given in T5, which coincides in the increase of acetic and butyric acids in caecal of broilers given dietary oregano aqueous extracts [109]. This finding, however does not confirm if the abundance of SCFA produced by gut microbes are optimal for gut health and performance in broilers, since exorbitant amounts of SCFAs may activate the gut microbiota-brain-cell axis response, resulting in either enteritis or other metabolic syndromes [110]. Furthermore, greater amounts of propionate and butyrate acids were previously detected in birds with low feed efficiency than those with high feed efficiency [111]. Nonetheless, a higher number of EPEF+ MAGs were shown to possess at least one KEGG module associated to SCFA production, compared to EPEF-MAGs, potentially indicating contribution of SCFA production capacity of caecal gut microbiota to broiler performance.

Our study boasts several strengths, including the commercial farm set-up representing real-life poultry industrial farming, the utilisation of shotgun metagenomic sequencing data, and thorough assessments of performances characteristics. These aspects empowered us to delve into the intricate composition and functions of gut microbiota concerning GHI administration with meticulous resolution and effective control of potential confounding factors. As our sampling was limited to a single genetic line of chickens and confined to caecal genetic study at one timepoint, we missed the opportunity to observe the effects of GHI on early development, and its potential links to temporal and spatial shifts of the chicken gut microbiome. For instance, a previous study by Gao et al., (2017) demonstrated that maturation of gut microbiota is promoted by probiotic administration whilst delayed by antibiotic use, highlighting the importance of broiler age in the use of supplements. Additional study into other gut compartments, timepoints, and other metagenomic features is therefore warranted. This includes deeper investigation of other gut microorganisms such as of bacteriophages and fungi, and of other relevant microbial elements including CRISPR-Cas systems, resistance, stress genes and virulence genes. In addition, future application of a multi-omics approach involving proteomics, meta transcriptomics, and metabolomics may confirm several of our hypotheses and uncover other areas we are not able to explore. Nevertheless, we believe our research represents a novel and comprehensive comparative investigation of the chicken metagenomic changes between ionophores and GHIs.

## Conclusion

Metagenomics has allowed us to gain insights on the bacterial population in the chicken caeca. This method has been utilised to highlight the differences in the composition, diversity, and metabolic functions of these caecal bacteria as influenced by different gut health schemes. Performing such analyses, we explored the structure of the gut bacterial population in conjunction with relevant metadata. We identified several MAGs such as *Rikenella microfusus,* UBA 7160 species and *Lactobacillus* species as beneficial organisms, due to their positive association with EPEF and higher capacity for metabolic functions. Such information will enhance our understanding of the highly complex relationship between gut microbes and optimal performance. It will also enable us to devise effective interventions and control strategies against enteric pathogens, which are important members of the poultry gut microbiome.

Among the gut health strategies investigated in this study, we observed that use of Probiotics B in a flock as observed in C2 enables better bird performance. Specifically, supplementation of Probiotics B in conjunction with vaccination is observed as the best GHI strategy, resulting in a similar performance to the control. However, we still observe the ionophore group to have the best performance and is hypothesised to be due to their ability to reduce microbial competition, resulting in a more efficient capture of nutrients by the gut microbiota and subsequently by the host.

Nonetheless, our results demonstrates that supplementation of GHIs are effective methods for broiler gut modulation, with evidence of having various influences on both MAG composition and feed related metabolic functions. Our data also suggests that excessive administration of GHIs may not be beneficial for performance, highlighting the importance of careful selection of GHI type and GHI combinations. These results significantly enhance our comprehension of microbiota-related metabolic pathways, offering new avenues to improve overall performance and poultry health.

## Declarations

### Availability of data and material

The sequencing datasets generated and/or analysed during the current study are available in the ENA repository Accession PRJEB75892.

## Authors’ contributions

Conceptualisation: AR, CH, UL

Laboratory work: GP, BS, CK

Data curation: GP, UI

Formal analysis: GP, UI, BS

Writing – original draft: GP

Writing – review and editing: GP, OG, AP, CK, CH, DX, AR, UL, CN, UI, BS

## List of abbreviations

ADG: Average daily gain
ANOVA: Analysis of variance
BW: Bird weight
CAZymes: Carbohydrate active enzymes
COM: Completeness (Genome)
CON: Contamination (Genome)
CP: Crude protein
DMRT: Duncan Multiple Range Test
EPEF: European poultry efficiency factor
F: Finisher diet
FA: Fisher’s alpha
FCR: Feed conversion ratio
FPD: Footpad dermatitis
G: Grower diet
GC: Guanosine-cytosine
GH: Glycoside hydrolase
GHI: Gut health intervention
GLLVM: General linear latent variable model
H: Shannon’s index
HB: Hockburn
KO: KEGG Orthology
LASSO: Least absolute shrinkage and selection operator
MAG: Metagenome-assembled genomes
MT: Mortality
NSP: Non-starch polysaccharides
PA: Probiotics A
PB: Probiotics B
PCA: Principal component analysis
PCoA: Principal coordinate analysis
PG: Phylogenetic gain (Novelty)
PERMANOVA: Permutational multivariate
ANOVA PL: Polysaccharide lyase
R: Richness
S: Specnumber
Si: Simpson
SCFA: Short-chain fatty acids
SCG: Single Copy Genes
TSS + CLR: Total Sum Scaling and Centralised Log Ratio
W: Withdrawal diet

## Acknowledgements

We are grateful to Abbie Graham and Hugo Hanna for their contribution in sampling and performance data collection. We also thank John Moore for his insights and comments on this study.

